# AMPK-dependent brain-wide norepinephrine release promotes recovery from metabolic coma in mice

**DOI:** 10.64898/2026.10.07.757341

**Authors:** Menghua Wu, Li Zhang, Yaning Li, Guoqing Chen, Dapeng Li, Miao Zhao, Ruoning Wang, Huating Gu, Huimeng Song, Ying Cao, Zhiwen Yang, Zhihui Liu, Fan Zhang, Tian-Zuo Li, Hui-Hui Miao, Peng Cao

## Abstract

Coma is a pathological brain state characterized by loss of consciousness. How the brain recovers from coma remains largely unknown. Using mouse models of metabolic coma, we identified an endogenous neuromodulatory pathway that actively promotes coma recovery. In mice subjected to hypoglycemic coma, hypoglycemia triggered AMPK-dependent excitation of nucleus tractus solitarius (NTS) neurons, which activated locus coeruleus (LC) norepinephrine neurons through a glutamatergic NTS-LC circuit. LC activation drives widespread norepinephrine release across cortical and subcortical regions, thereby promoting post-coma arousal. Notably, coma-associated brain-wide norepinephrine release was unconventional in both scale and duration, with a magnitude approximately 10-fold higher than physiological norepinephrine release during sleep-wake cycles. This neuromodulatory process also operated during hypercapnic coma. Vagus nerve stimulation accelerated coma recovery by engaging this endogenous anti-coma mechanism. Our findings uncover an AMPK-dependent brainstem circuit that actively restores arousal from metabolic coma, revealing a neural defense strategy against profound metabolic stress.

## Introduction

Coma is a pathological brain state in which consciousness is profoundly impaired^1,2^. This condition arises from diverse etiologies, including metabolic encephalopathies such as hypoglycemia and hypercapnia, traumatic brain injury, intracerebral hemorrhage, and other acute neurological insults^3^. Despite substantial advances in clinical neurocritical care, the recovery of consciousness following coma remains a key unresolved challenge in neuroscience^4^.

In humans, electroencephalography (EEG)^5,6^ and neuroimaging^7,8^ have provided critical insights into brain network failure during coma. These studies indicate that coma involves disrupted functional connectivity between the corticothalamic network and neuromodulatory systems originating from the brainstem, hypothalamus, and basal forebrain^9^. Among these systems, the ascending reticular activating system (ARAS) in the brainstem occupies a central position in arousal regulation and has been strongly implicated in disorders of consciousness^10,11^. Therapeutic approaches that engage neuromodulatory circuits, including deep brain stimulation^12^, vagus nerve stimulation (VNS)^13,14^, and pharmacological treatments^15^, have therefore attracted increasing interest as strategies to promote recovery of consciousness.

Mechanistic studies in humans are limited by ethical constraints and restricted accessibility of relevant neural circuits. Animal models have consequently become essential for exploring the neurobiological mechanisms that govern emergence from coma^16–19^. Classical electrical stimulation and lesion studies have established a causal link between ARAS nuclei and arousal control^20,21^. However, the ARAS contains dozens of nuclei composed of molecularly and functionally distinct neuronal subtypes, each capable of shaping corticothalamic activity through the release of different neurotransmitters. Given this complexity, the mechanisms through which defined neuronal populations detect coma-associated pathological stress and initiate arousal-promoting responses remain largely unresolved.

Mice provide a powerful system for dissecting the molecular and cellular mechanisms underlying recovery of consciousness following coma, supported by extensive genetic toolkits for precise circuit and molecular interrogation^22^. Nevertheless, standardized methods for inducing and quantifying coma-like states are currently lacking. In this study, we developed two experimental paradigms to model metabolic coma and post-coma arousal in mice. Using these paradigms, we interrogated how defined neuromodulatory systems contribute to the restoration of consciousness from coma.

## Results

### Experimental paradigm for studying hypoglycemic coma in mice

Excess insulin administration can precipitate hypoglycemic coma in patients with diabetes, and insulin similarly induces hypoglycemic coma in mice^23^. We therefore used mice to develop a paradigm for induction and quantification of insulin-induced hypoglycemic coma (<u>Supplementary Video 1</u>). After 24 h of fasting, mice received an intraperitoneal injection of insulin (12 U/kg), which produced rapid but reversible hypoglycemic symptoms, including reductions in blood glucose level, respiratory rate, heart rate, and body temperature (Extended Data Fig. 1a–c). Approximately 1.2 h after insulin injection (<u>latency = 1.2 ± 0.3 h</u>), all mice (<u>15/15</u>) developed loss of righting reflex (LORR) (Fig. 1a), a behavioral state that persisted for about 3 h (<u>duration = 3.1 ± 0.4 h</u>). During LORR, mice showed a weaker pupillary light reflex than during wakefulness (Extended Data Fig. 1d, e), consistent with clinical features observed in comatose patients^24^.

**Fig. 1.**
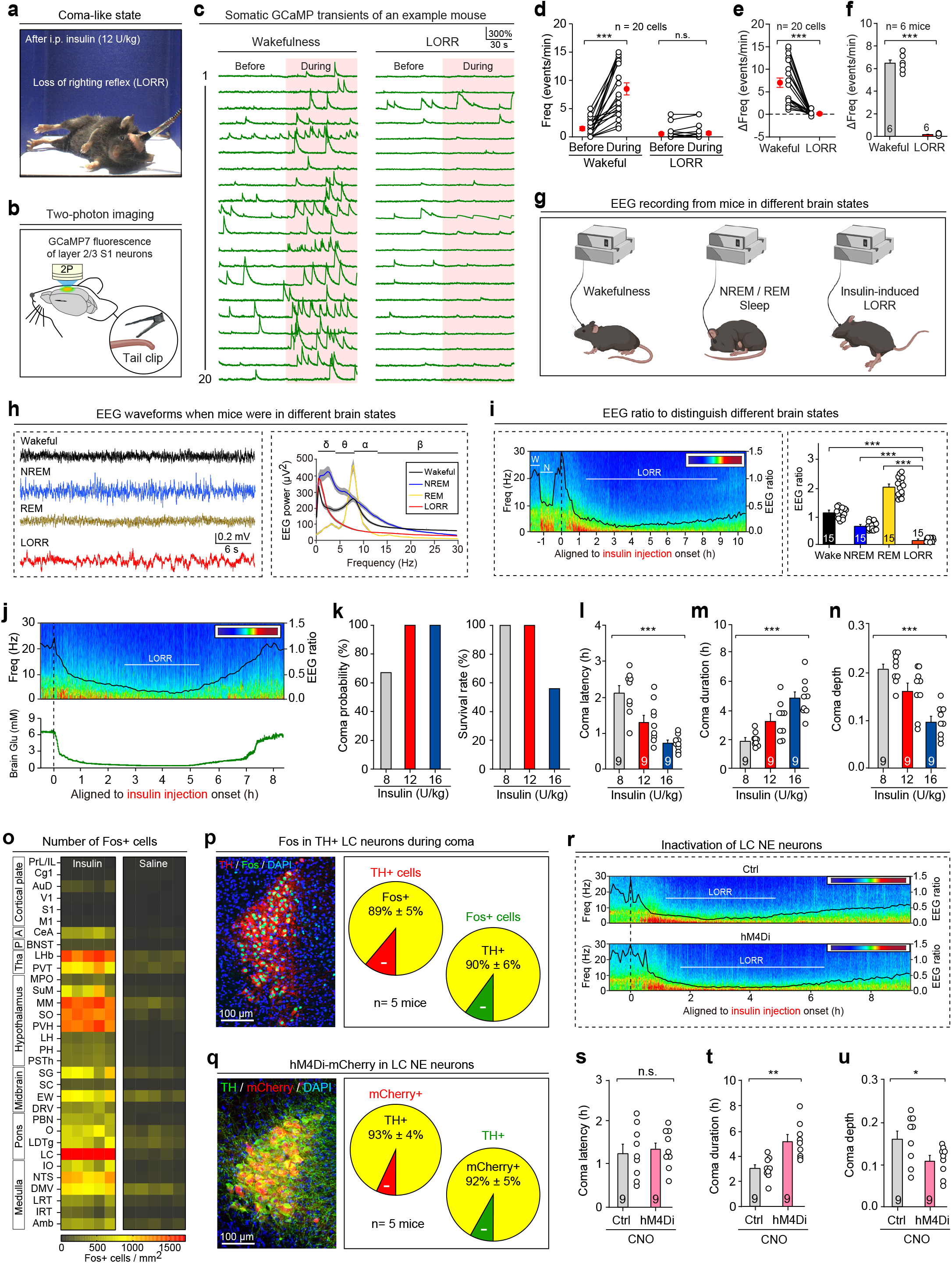
Insulin-induced hypoglycemic coma and involvement of LC NE neurons. **(a)** Representative image showing loss of righting reflex (LORR) in a mouse after insulin treatment. **(b)** Schematic showing two-photon imaging of layer 2/3 cortical neurons in the S1 of mice during noxious mechanical stimulation on the tail. **(c)** GCaMP traces from layer 2/3 S1 cortical neurons of a representative mouse during wakefulness and LORR, showing somatic calcium transients (SCTs) in response to noxious mechanical stimulation. **(d)** Quantification of SCT frequency before and during mechanical stimulus of a representative mouse in wakefulness and LORR. **(e, f)** Quantification of ΔFreq in a representative mouse (e) and across multiple mice (f) in wakefulness and LORR. **(g)** Schematic showing EEG recording in mice during wakefulness, NREM/REM sleep, and LORR. **(h)** Representative EEG traces (*left*) and EEG power distributions across frequencies during wakefulness, NREM/REM sleep, and LORR (*right*). **(i)** EEG heatmap, EEG ratio curve, and LORR timing in a representative mouse (*left*), and quantification of EEG ratios during wakefulness, NREM/REM sleep, and LORR across 15 mice (*right*). W, wakefulness. N, NREM sleep. **(j)** EEG heatmap, EEG ratio curve, and LORR timing in a representative mouse (*top*), and cortical glucose level before and after insulin injection (*bottom*). Vertical dashed line indicates time point of insulin injection. **(k)** Quantification of coma probability (*left*) and survival rate (*right*) in mice treated with different insulin doses. **(l-n)** Quantification of coma latency (l), coma duration (m), and coma depth (n) in mice treated with different insulin doses. **(o)** Quantification of Fos+ cell density in different brain regions in five pairs of mice. Brain-section images are shown in Figure S2B. **(p)** Representative micrograph (*left*) and quantification (*right*) of Fos expression in TH+ LC NE neurons. **(q)** Representative micrograph (*left*) and quantification (*right*) of hM4Di-mCherry expression specifically in TH+ LC NE neurons. **(r)** EEG heatmap, EEG ratio curve, and LORR timing showing insulin-induced coma in mice with or without chemogenetic inactivation of LC NE neurons. **(s-u)** Quantification of coma latency (s), coma duration (t), and coma depth (u) in mice with or without chemogenetic inactivation of LC NE neurons. Scale bars are labeled in graphs. Numbers of cells (d, e) or mice (f, i, l-n, p, q, s-u) are indicated in graphs. Data in (d-f, i, l-n, p, q, s-u) are mean ± SEM. Statistical analyses (d-f, i, l-n, s-u) were performed using Student *t-*test (*** *P* < 0.001; ** *P* < 0.01; * *P* < 0.05; n.s. *P* > 0.1) or one-way ANOVA (*** *P* < 0.001). For *P*-values, see Supplementary Table 3.

To determine whether insulin-induced LORR reflects a coma-like state, we assessed cortical responsiveness to noxious stimuli, as markedly reduced cortical responses to painful stimuli represent a hallmark of coma in humans^24^. AAV-hSyn-GCaMP7 was injected into the primary somatosensory cortex (S1), resulting in GCaMP7 expression in S1 neurons (Extended Data Fig. 1f). Two-photon imaging was then used to record GCaMP signals from layer 2/3 S1 neurons before and during a noxious mechanical stimulus to the tail with an alligator clip (250 g, 2 min) (Fig. 1b). The episodes of somatic calcium transients (SCTs) were compared in the same S1 neurons during wakefulness and LORR (Fig. 1c). In awake mice, noxious stimulation evoked a robust increase in SCT frequency; in contrast, the same stimulus failed to increase SCT frequency during LORR (Fig. 1d). We defined ΔFreq as the change in SCT frequency after stimulus application. During LORR, ΔFreq remained close to zero in both representative cells from an individual mouse (Fig. 1e) and pooled data from multiple mice (Fig. 1f). These results suggest a loss of responsiveness in S1 neurons to noxious stimuli during LORR, supporting the hypothesis that insulin-induced LORR represents a coma-like state in mice. Accordingly, LORR latency and duration were used to define the latency and duration of the coma-like state in mice.

To quantify coma depth, EEG activity was recorded in mice across wakefulness, NREM/REM sleep, and insulin-induced LORR (Fig. 1g). EEG waveforms (Fig. 1h<u>, *left*</u>) and spectral power distribution (Fig. 1h<u>, *right*</u>) during LORR differed markedly from those recorded during other brain states. We defined the EEG ratio as the average power in theta (θ) band divided by that in delta (δ) band. The EEG ratio fluctuated as mice transitioned through wakefulness (W), NREM sleep (N), and insulin-induced LORR (Fig. 1i<u>, *left*</u>). Across 15 mice, the minimum EEG ratio during LORR was significantly lower than the values observed in all other brain states (Fig. 1i<u>, *right*</u>). After insulin treatment, the EEG ratio progressively declined, reached a minimum, and then recovered, forming a U-shaped trajectory that paralleled changes in blood glucose level (Extended Data Fig. 1g), respiratory and heart rates (Extended Data Fig. 1h), body temperature (Extended Data Fig. 1i), and intracerebral glucose level (Extended Data Fig. 1j <u>and</u> Fig. 1j). Based on these observations, the minimum EEG ratio during LORR was used as a quantitative index of coma-like state depth.

We next examined the relationship between insulin dose and coma-like state. Three doses of insulin (8, 12, and 16 U/kg) were tested, each of which induced marked hypoglycemia (Extended Data Fig. 1k). Both recovery time and the minimum blood glucose level after insulin injection varied in a dose-dependent manner (Extended Data Fig. 1l, m). The proportion of mice developing a coma-like state approached 100% after administration of 12 or 16 U/kg insulin (Fig. 1k<u>, *left*</u>). However, survival after insulin injection decreased to approximately 60% at 16 U/kg (Fig. 1k<u>, *right*</u>), likely attributable to severe hypoglycemia-induced myoclonic seizures^23^. These data suggest that 12 U/kg insulin provides a safe and efficient dose for coma induction. Comparison of coma-like state parameters across insulin doses further showed that increasing insulin dose shortened coma latency (Fig. 1l), prolonged coma duration (Fig. 1m), and increased coma depth (Fig. 1n). Together, these results establish 12 U/kg insulin as a reliable and safe dose for inducing hypoglycemic coma in mice.

### Locus coeruleus (LC) norepinephrine (NE) neurons counteract hypoglycemic coma

We next mapped brain areas that were activated during insulin-induced hypoglycemic coma by Fos immunostaining. Wild-type mice fasted for 24 h received an intraperitoneal injection of insulin (12 U/kg) to induce hypoglycemic coma. After recovery from coma, mice were perfused and processed for subsequent histological analysis (Extended Data Fig. 2a). Brain sections from saline-treated control mice and insulin-treated mice were immunostained with an anti-Fos antibody, which revealed Fos expression across multiple brain regions (Extended Data Fig. 2b). Quantitative analysis showed selective enrichment of Fos+ cells in subcortical regions of comatose mice (Fig. 1o). Among these regions, the LC contained abundant Fos+ cells, approximately 90% of which were tyrosine hydroxylase (TH)+ NE neurons (Fig. 1p <u>and</u> Extended Data Fig. 2c).

To determine whether LC NE neurons contribute to hypoglycemic coma, AAV-DIO-hM4Di-mCherry or control AAV-DIO-mCherry was bilaterally injected into the LC of Dbh-Cre mice (Extended Data Fig. 2d, e). The expression of hM4Di-mCherry was restricted to TH+ LC NE neurons (Fig. 1q). In acute brain slices containing the LC, CNO effectively suppressed action potential firing in hM4Di-expressing LC NE neurons (LC-NE^hM4Di^ neurons) (Extended Data Fig. 2f, g). Both LC-NE^hM4Di^ and control mice were pretreated with CNO (1 mg/kg) 20 min before insulin injection, and LORR behavior and cortical EEG were recorded simultaneously to quantify the latency, duration, and depth of hypoglycemic coma (Fig. 1r). Chemogenetic inactivation of LC-NE^hM4Di^ neurons did not alter coma latency but significantly prolonged coma duration (Fig. 1s, t). EEG analysis further showed that inhibition of LC NE neurons increased coma depth, as indicated by lower minimum values of the EEG ratio in LC-NE^hM4Di^ mice compared with control mice (Fig. 1u). These findings suggest that LC NE neuron activity opposes insulin-induced hypoglycemic coma and facilitates recovery.

### Brain-wide NE release during hypoglycemic coma

We next investigated how LC NE neurons counteract hypoglycemic coma. Because LC NE neurons innervate broad brain regions and modulate brain states through NE release^25,26^, we monitored cortex-wide NE dynamics by mesoscopic imaging of neuronal GRAB-NE2m signals in Snap25-Cre::RCL-jRGECO1a-2A-NE2m mice (Fig. 2a, b). A 470-nm laser was used to excite GRAB-NE2m, and a 405-nm laser was used for correction of cortical hemodynamic signals^27^. Following insulin injection, GRAB-NE2m fluorescence signals progressively increased across multiple cortical regions, including the primary motor cortex (M1), S1, and the primary visual cortex (V1) (Fig. 2c, d), suggesting broad cortical NE release during hypoglycemia.

**Fig. 2.**
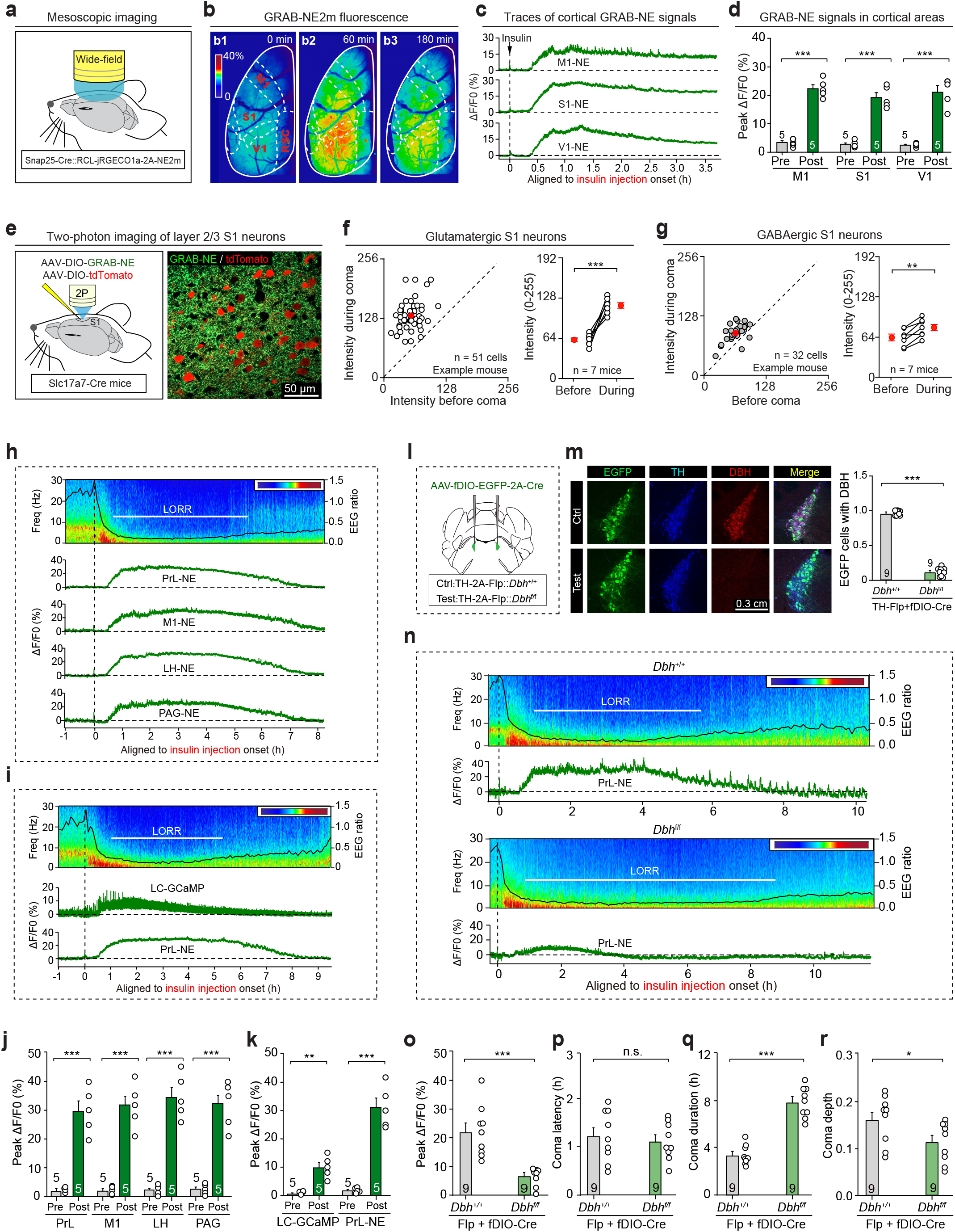
Hypoglycemia-induced brain-wide NE release is critical for recovery from coma. (a, **b)** Schematic (a) and representative images (b) showing mesoscopic imaging of GRAB-NE2m signals in cortical areas of mice after insulin treatment. b1–b3 show heatmaps of cortical GRAB-NE2m fluorescence after insulin injection. **(c, d)** Representative traces (c) and quantification (d) of peak ΔF/F0 of GRAB-NE2m signals in different cortical areas (M1, S1, and V1) before and after insulin treatment. **(e)** Schematic (*left*) showing two-photon imaging of GRAB-NE2m signals in individual glutamatergic S1 neurons (*right*). **(f)** Comparison of GRAB-NE2m fluorescence intensity before and during insulin-induced coma in individual glutamatergic S1 neurons from a representative mouse (*left*) and across multiple mice (*right*). **(g)** Comparison of GRAB-NE2m fluorescence intensity before and during insulin-induced coma in individual GABAergic S1 neurons in a representative mouse (*left*) and across multiple mice (*right*). **(h)** Representative traces of GRAB-NE2m signals in cortical (PrL, M1) and subcortical regions (LH, PAG), shown with hypoglycemia-induced coma state. **(i)** Representative traces of GCaMP signals in LC NE neurons and GRAB-NE2m signals in the PrL, shown with hypoglycemia-induced coma state. **(j)** Quantification of peak ΔF/F0 of GRAB-NE2m signals in cortical (PrL, M1) and subcortical regions (LH, PAG) before and after insulin injection. **(k)** Quantification of peak ΔF/F0 of GCaMP signals in LC NE neurons and GRAB-NE2m signals in the PrL before and after insulin injection. **(l)** Schematic showing the strategy used to delete *Dbh* in LC NE neurons. **(m)** Representative micrographs (*left*) and quantification (*right*) showing efficient *Dbh* deletion in LC NE neurons. **(n)** Representative traces of GRAB-NE2m signals in the PrL, shown with hypoglycemia-induced coma state in mice with or without *Dbh* deletion in LC NE neurons. **(o)** Quantification showing effects of *Dbh* deletion on peak ΔF/F0 of GRAB-NE2m signals in the PrL. **(p-r)** Quantification showing effects of *Dbh* deletion in LC NE neurons on hypoglycemic coma latency (p), duration (q), and depth (r). Scale bars are labeled in graphs. Numbers of neurons (f, g) or mice (d, f, g, j, k, m, o-r) are indicated in graphs. Data in (d, f, g, j, k, m, o-r) are mean ± SEM. Statistical analyses (d, f, g, j, k, m, o-r) were performed using Student *t-*test (*** *P* < 0.001; ** *P* < 0.01; * *P* < 0.05; n.s. *P* > 0.1). For *P*-values, see Supplementary Table 3.

To resolve hypoglycemia-induced NE release at cellular resolution, we performed two-photon imaging of GRAB-NE2m signals in defined S1 neuronal populations. AAV-DIO-GRAB-NE2m and AAV-DIO-tdTomato were co-injected into S1 of Slc17a7-Cre mice, resulting in GRAB-NE2m expression in glutamatergic S1 neurons (Extended Data Fig. 2h, i). Imaging of layer 2/3 S1 neurons revealed a pronounced increase in GRAB-NE2m fluorescence 2 h after insulin injection, when mice had entered the coma-like state (Fig. 2e, f). Using the similar strategy, we expressed GRAB-NE2m and tdTomato in S1 GABAergic interneurons (Extended Data Fig. 2j, k). GRAB-NE2m fluorescence in layer 2/3 GABAergic interneurons modestly but significantly increased after entry into the coma-like state (Fig. 2g). These cellular-resolution measurements confirmed that insulin-induced hypoglycemic coma is accompanied by local NE release in cortical circuits.

We then used fiber photometry to determine whether hypoglycemia evokes NE release beyond the cortex. AAV-hSyn-GRAB-NE2m was ipsilaterally injected into multiple cortical and subcortical brain areas in the same mice, including the prelimbic cortex (PrL), the M1, the lateral hypothalamic area (LH), and the periaqueductal gray (PAG), followed by optical fiber implantation above each target region (Extended Data Fig. 3a, b). GRAB-NE2m signals were recorded with alternating 470- and 405-nm excitation, with 470-nm laser for GRAB-NE2m excitation and 405-nm laser for correction of movement artifacts, photobleaching and pH changes. Insulin injection elicited robust increases in GRAB-NE2m signals across these regions (Fig. 2h, j). Bilateral cortical recordings further showed that NE release was largely symmetrical between the left and right hemispheres (Extended Data Fig. 3c-f). Simultaneous recordings of PrL GRAB-NE2m signals and GCaMP7 activity in LC NE neurons (Extended Data Fig. 3g, h) revealed synchronized dynamics between NE release and activation of LC NE neurons (Fig. 2i, k). As a control, insulin injection did not alter GRAB-NEmut signals in the PrL, confirming the specificity of GRAB-NE2m signals (Extended Data Fig. 3i, j). Together, these results demonstrate that hypoglycemia induces widespread NE release across cortical and subcortical brain regions.

Using PrL GRAB-NE2m signals as a proxy, we next characterized the temporal profile of hypoglycemia-induced NE release. The rise time of GRAB-NE2m signal was defined as the time at which the signal reached 15% of peak amplitude relative to baseline. After insulin injection, PrL GRAB-NE2m signals began to rise at approximately 38 min (38 ± 7 min, *n* = 9 mice) and reached peak amplitude at approximately 107 min (107 ± 13 min, *n* = 9 mice). Compared with coma onset, GRAB-NE2m signals started to rise approximately 36 min earlier (36 ± 11 min, *n* = 9 mice). Simultaneous monitoring of GRAB-NE2m signals and blood glucose level showed that NE release began when blood glucose fell below 2.2 mM (2.2 ± 0.1 mM, *n* = 9 mice), whereas LORR emerged when blood glucose level declined below 1.1 mM (1.1 ± 0.1 mM, *n* = 9 mice). These data indicate that cortical NE release precedes coma onset.

To assess the magnitude of hypoglycemia-induced NE release, we performed two comparative analyses. First, hypoglycemia-induced GRAB-NE2m responses were compared with NE dynamics across natural sleep-wake transitions in the same mice. Notably, GRAB-NE2m signals increased during the wake-to-coma transition (Extended Data Fig. 3k) and decreased during the wake-to-sleep transition (Extended Data Fig. 3l), revealing opposing trends between the two transitions. In the same mice (*n* = 6), the hypoglycemia-induced GRAB-NE2m response (24.8% ± 4.3%) was approximately 10-fold larger than the response to the wake-to-NREM transition (2.4% ± 0.4%) and approximately 4-fold larger than the response to the wake-to-REM transition (6.1% ± 1.8%) (Extended Data Fig. 3m). These findings indicate that hypoglycemia evokes a unique NE response that is substantially larger than physiological NE fluctuations associated with sleep-wake cycling. Second, hypoglycemia-induced NE release was compared with the release of other neuromodulators, such as dopamine (Extended Data Fig. 3n, o). Interestingly, the hypoglycemia-induced GRAB-NE2m signal was significantly larger than the corresponding GRAB-DA2m signal (Extended Data Fig. 3p). These results suggest that hypoglycemia-induced NE release, rather than dopamine release, represents a prominent neuromodulatory signature of insulin-induced hypoglycemic coma.

### Critical role of NE release in counteracting hypoglycemic coma

We next investigated the functional contribution of brain-wide NE release to insulin-induced hypoglycemic coma. Dopamine β-hydroxylase (Dbh) is the rate-limiting enzyme for NE synthesis. To suppress NE release, *Dbh* was selectively deleted in LC NE neurons (Fig. 2l, m). *Dbh* deletion effectively abolished hypoglycemia-induced NE release in the PrL (Fig. 2n, o). This genetic manipulation did not alter the latency of insulin-induced hypoglycemic coma (Fig. 2p), but significantly prolonged coma duration (Fig. 2q) and increased coma depth (Fig. 2r). As a control, *Gal*, which encodes the neuropeptide galanin, was deleted selectively in LC NE neurons (Extended Data Fig. 4a, b). *Gal* deletion did not alter hypoglycemia-induced cortical NE release (Extended Data Fig. 4c, d), and had no effect on coma latency, duration, or depth (Extended Data Fig. 4e-g). These findings indicate that widespread NE release from LC neurons may play a critical role in limiting the duration and severity of hypoglycemic coma.

### NTS-LC circuit drives widespread NE release

We next sought to identify the upstream circuits that couple hypoglycemia to widespread NE release in the brain. Recombinant G-deleted rabies virus (RV) tracing was used to map monosynaptic inputs to LC NE neurons (Extended Data Fig. 5a, b)^28,29^. RV-labeled neurons were detected in the nucleus tractus solitarius (NTS) and several other brain regions (Extended Data Fig. 5c, d). To validate the NTS-LC projection, AAV-hSyn-ChR2-mCherry was injected into the NTS of TH-GFP mice, which labeled ChR2-mCherry+ axons within the LC (Extended Data Fig. 5e, f). In acute brain slices, photostimulation of ChR2-mCherry+ axons evoked postsynaptic currents in TH-GFP+ LC neurons (Extended Data Fig. 5g, h). These currents were insensitive to the GABAa receptor antagonist picrotoxin (PTX) but were abolished by the glutamate receptor antagonists D-AP5 and CNQX, indicating glutamatergic transmission through the NTS-LC circuit (Extended Data Fig. 5h, i). Together, these anatomical and electrophysiological data demonstrate that a subset of NTS neurons forms a glutamatergic circuit to monosynaptically innervate LC NE neurons.

We subsequently asked whether insulin-induced hypoglycemia recruits this NTS-LC circuit. AAV2-retro-Cre was injected into the LC and AAV-DIO-tdTomato was injected into the NTS of wild-type mice to label LC-projecting NTS neurons (Fig. 3a, b). Following insulin-induced hypoglycemic coma, nearly 70% of tdTomato+ neurons expressed Fos (73% ± 6%, *n* = 3 mice; Fig. 3c), suggesting strong activation of LC-projecting NTS neurons during hypoglycemia. To monitor this activation together with cortical NE release, AAV2-retro-Cre, AAV-DIO-GCaMP7, and AAV-GRAB-NE2m were injected into the LC, NTS, and PrL, respectively, followed by optical fiber implantation above the NTS and PrL (Fig. 3d <u>and</u> Extended Data Fig. 5j, k). Simultaneous recordings showed that GCaMP signals in LC-projecting NTS neurons rose and declined in parallel with PrL GRAB-NE2m signals (Fig. 3e, f). These findings indicate that hypoglycemia activates LC-projecting NTS neurons in close temporal association with cortical NE release.

**Fig. 3.**
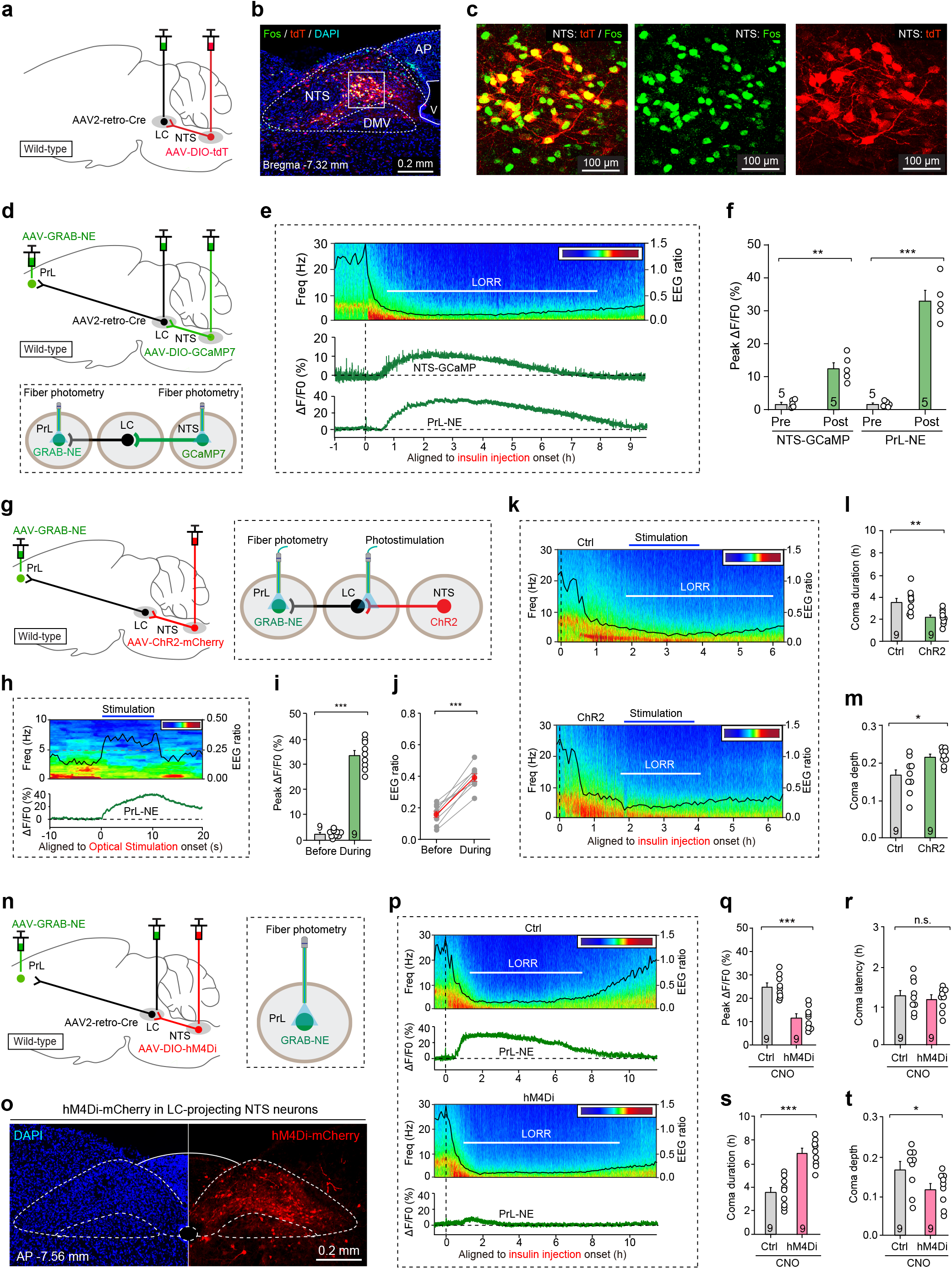
LC-projecting NTS neurons drive cortical NE release during hypoglycemic coma. **(a)** Schematic showing strategy to label LC-projecting NTS neurons in wild-type mice. **(b, c)** Representative coronal brain section (b) and magnified fields (c) showing that tdTomato-labeled LC-projecting NTS neurons were mostly positive for hypoglycemia-induced Fos expression. **(d)** Schematic showing strategy used for simultaneous recording of GCaMP signals in LC-projecting NTS neurons and GRAB-NE2m signals in the PrL. **(e)** Representative traces of GCaMP signals in LC-projecting NTS neurons and GRAB-NE2m signals in the PrL, shown with hypoglycemic coma state. **(f)** Quantification of peak ΔF/F0 of GCaMP signals in LC-projecting NTS neurons and GRAB-NE2m signals in the PrL before and after insulin injection. **(g)** Schematic showing strategy used to measure NE release in the PrL evoked by photostimulation of ChR2-mCherry+ NTS-LC axon terminals. **(h)** Representative EEG heatmap, EEG ratio curve and GRAB-NE2m signal before, during, and after photostimulation of NTS-LC circuit. **(i)** Quantification of peak ΔF/F0 of GRAB-NE2m signals in PrL before and during photostimulation of NTS-LC circuit. **(j)** Quantification of EEG ratio before and during photostimulation of NTS-LC circuit. **(k)** Representative EEG heatmaps, EEG ratio curves, and LORR timing showing insulin-induced coma in mice with or without repetitive photostimulation of NTS-LC circuit. **(l, m)** Quantification of coma duration (l) and coma depth (m) showing effect of repetitive photostimulation of NTS-LC circuit on hypoglycemia-induced coma. **(n)** Schematic showing strategy used for specific expression of hM4Di-mCherry in LC-projecting NTS neurons and fiber recordings of GRAB-NE2m signals in the PrL. **(o)** Representative micrograph showing expression of hM4Di-mCherry in LC-projecting NTS neurons. **(p)** Representative traces of GRAB-NE2m signals in the PrL, shown with hypoglycemia-induced coma state in mice with or without chemogenetic inactivation of LC-projecting NTS neurons. **(q)** Quantification of peak GRAB-NE2m ΔF/F0, showing effect of chemogenetic inactivation of LC-projecting NTS neurons on hypoglycemia-induced NE release in the PrL. **(r-t)** Quantification of coma latency (r), duration (s), and depth (t), showing effect of chemogenetic inactivation of LC-projecting NTS neurons on insulin-induced hypoglycemic coma. Scale bars are labeled in graphs. Numbers of mice (f, i, j, l, m, q-t) are indicated in graphs. Data in (f, i, j, l, m, q-t) are mean ± SEM. Statistical analyses (f, i, j, l, m, q-t) were performed using Student *t-*test (*** *P* < 0.001; ** *P* < 0.01; * *P* < 0.05; n.s. *P* > 0.1). For *P*-values, see Supplementary Table 3.

We next determined whether activation of the NTS-LC projection is sufficient to drive cortical NE release and alter hypoglycemic coma. AAV-hSyn-ChR2-mCherry and AAV-hSyn-GRAB-NE2m were injected into NTS and PrL, respectively, followed by optical fiber implantation above the LC and PrL (Fig. 3g). In comatose mice, brief photostimulation of ChR2-mCherry+ axons in the LC (10 s, 20 Hz, 5 mW) evoked robust NE release in the PrL (Fig. 3h, i), accompanied by a significant increase in the EEG ratio (Fig. 3h, j). These effects occurred without detectable changes in blood glucose level (Extended Data Fig. 5l, m), indicating that NTS-LC circuit modulates coma-associated arousal without directly altering systemic glucose levels under these conditions. When mice entered hypoglycemic coma, repeated photostimulation with the same parameters once per minute for 2 h significantly shortened coma duration and reduced coma depth (Fig. 3k-m).

Conversely, we examined whether chemogenetic inactivation of the NTS-LC circuit attenuates cortical NE release and worsens hypoglycemic coma. AAV-hSyn-GRAB-NE2m, AAV2-retro-Cre, and AAV-DIO-hM4Di-mCherry were injected into the PrL, LC, and NTS, respectively, followed by optical fiber implantation above the PrL (Fig. 3n). This strategy drove hM4Di-mCherry expression in LC-projecting NTS neurons (Fig. 3o). Chemogenetic inhibition of these neurons with CNO (1 mg/kg) significantly reduced insulin-evoked cortical NE release (Fig. 3p, q). Silencing these neurons did not alter coma latency but markedly prolonged coma duration and increased coma depth (Fig. 3r-t). Collectively, these results identify the NTS-LC projection as an upstream glutamatergic circuit that drives widespread NE release and counteracts hypoglycemic coma.

### Hypoglycemia activates LC-projecting NTS neurons via AMPK

We next investigated how hypoglycemia activates LC-projecting NTS neurons. In acute brain slices, we performed current-clamp recordings from LC-projecting NTS neurons, measuring their responses to low-glucose (1 mM) ACSF treatment (Fig. 4a). Quantification of action potential firing and membrane potential changes indicated that most recorded neurons (18/21, 86%) responded to low glucose with increased firing and membrane depolarization (Fig. 4b, c). These results indicate that most LC-projecting NTS neurons directly sense reduced glucose availability and exhibit glucose-inhibited (GI) properties.

**Fig. 4.**
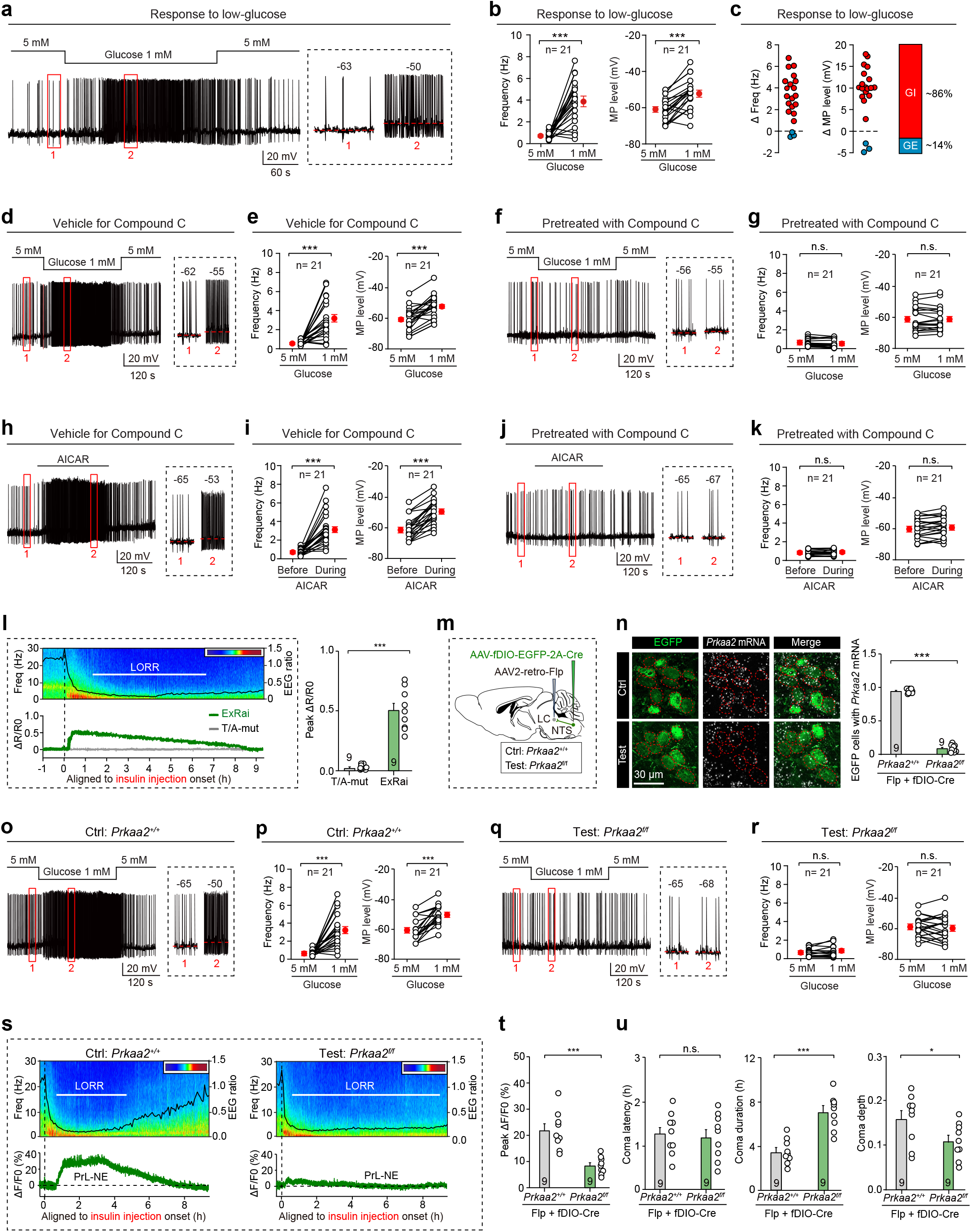
Hypoglycemia-induced brain-wide NE release depends on AMPK signaling in LC-projecting NTS neurons. **(a)** Representative traces of action potential firing (*left*) and membrane potential (*right*) showing responses of an LC-projecting NTS neuron to low-glucose stimulation. **(b)** Quantification of action potential firing frequency (*left*) and membrane potential level (*right*) in all recorded LC-projecting NTS neurons to low-glucose stimulation. **(c)** Quantification of changes in firing frequency (ΔFreq) and membrane potential (ΔMP), showing that approximately 86% of neurons exhibited glucose-inhibited (GI) properties. **(d, f)** Representative traces of action potential firing (*left*) and membrane potential (*right*) showing low-glucose responses of LC-projecting NTS neurons in the absence (d) or presence (f) of Compound C perfusion. **(e, g)** Quantification of firing frequency (*left*) and membrane potential level (*right*) in all recorded neurons to low-glucose stimulation in the absence (e) or presence (g) of Compound C perfusion. **(h, j)** Representative traces of action potential firing (*left*) and membrane potential (*right*) in LC-projecting NTS neurons in response to AICAR in the absence (h) or presence (j) of Compound C perfusion. **(i, k)** Quantification of firing frequency (*left*) and membrane potential level (*right*) in all recorded neurons in response to AICAR in the absence (i) or presence (k) of Compound C perfusion. **(l)** Representative traces (*left*) and quantification (*right*) of ExRai AMPKAR 470/405 ratio changes, showing an increase in AMPK activity in LC-projecting NTS neurons after insulin injection. T/A-mut was used as a negative control. **(m)** Schematic showing strategy used to delete *Prkaa2* specifically in LC-projecting NTS neurons. **(n)** Representative micrographs (*left*) and quantification (*right*) of EGFP+ cells containing *Prkaa2* mRNA, showing effective *Prkaa2* deletion in LC-projecting NTS neurons. **(o, q)** Representative traces of action potential firing (*left*) and membrane potential (*right*) in response to low-glucose stimulation in LC-projecting NTS neurons without (o) or with (q) *Prkaa2* deletion. **(p, r)** Quantification of firing frequency (*left*) and membrane potential level (*right*) in response to low-glucose stimulation in LC-projecting NTS neurons without (p) or with (r) *Prkaa2* deletion. **(s)** Representative traces of GRAB-NE2m signals (*bottom*), shown with hypoglycemia-induced coma state (*top*) in mice with or without *Prkaa2* deletion in LC-projecting NTS neurons. **(t)** Quantification of peak GRAB-NE2m ΔF/F0 in the PrL of mice with or without *Prkaa2* deletion in LC-projecting NTS neurons. **(u)** Quantification showing effects of *Prkaa2* deletion in LC-projecting NTS neurons on insulin-induced hypoglycemic coma latency (*left*), duration (*middle*), and depth (*right*). Scale bars are labeled in graphs. Numbers of neurons (b, e, g, i, k, p, r) or mice (l, n, t, u) are indicated in graphs. Data in (b, e, g, i, k, l, n, p, r, t, u) are mean ± SEM. Statistical analyses (b, e, g, i, k, l, n, p, r, t, u) were performed using Student *t-*test (*** *P* < 0.001; * *P* < 0.05; n.s. *P* > 0.05). For P values, see Supplementary Table 3.

AMP-activated protein kinase (AMPK) has been implicated as a key molecular sensor of cellular energy deficiency in GI neurons^30,31^. To determine whether AMPK participates in glucose sensing in LC-projecting NTS neurons, we tested the effect of Compound C, a selective AMPK inhibitor, on low-glucose-evoked excitation. Vehicle (DMSO)-treated LC-projecting NTS neurons showed robust increases in action potential firing and membrane depolarization during low-glucose stimulation (Fig. 4d, e). In contrast, pretreatment with Compound C (10 μM) prevented both the firing increase and depolarization induced by low glucose (Fig. 4f, g). Conversely, perfusion with ACSF containing the AMPK agonist AICAR (10 μM) activated LC-projecting NTS neurons (Fig. 4h, i), an effect that was blocked by Compound C pretreatment (Fig. 4j, k). These pharmacological data suggest that AMPK activity may contribute to glucose sensing in LC-projecting NTS neurons.

We subsequently investigated whether AMPK activity in LC-projecting NTS neurons contributes to hypoglycemia-induced NE release *in vivo*. Brain ATP levels were markedly reduced 1 h after insulin injection (Extended Data Fig. 6a). To determine whether this energy deficit increased AMPK activity in LC-projecting NTS neurons, we expressed ExRai AMPKAR, a reporter of AMPK activity^32^, in these neurons and monitored AMPK activity by fiber photometry (Extended Data Fig. 6b). Insulin treatment rapidly increased the AMPKAR 470/405 ratio in LC-projecting NTS neurons, whereas no ratio changes were observed in neurons expressing T/A-mutated AMPKAR as a control (Fig. 4l). These results demonstrate that hypoglycemia elevates AMPK activity in LC-projecting NTS neurons *in vivo*.

We next asked whether AMPK activation in LC-projecting NTS neurons is required for cortical NE release and coma resistance during hypoglycemia. Neurons constitutively express *Prkaa1* and *Prkaa2*, which encode catalytic subunits AMPKα1 and AMPKα2, respectively^33^. AAV2-retro-Flp was injected into the LC, and AAV-fDIO-EGFP-2A-Cre was injected into the NTS of *Prkaa2*^f/f^ mice (Fig. 4m), resulting in effective *Prkaa2* deletion in LC-projecting NTS neurons (Fig. 4n). In control mice without *Prkaa2* deletion, LC-projecting NTS neurons retained strong excitatory responses to low-glucose stimulation (Fig. 4o, p). In contrast, *Prkaa2* deletion abolished low-glucose responsiveness in these neurons (Fig. 4q, r). *Prkaa2* deletion also significantly suppressed hypoglycemia-induced cortical NE release (Fig. 4s, t). Although coma latency remained unchanged, coma duration and depth were markedly increased (Fig. 4u). In a parallel set of experiments, *Prkaa1* was deleted in LC-projecting NTS neurons (Extended Data Fig. 6c, d). Unlike *Prkaa2* deletion, *Prkaa1* deletion in LC-projecting NTS neurons failed to abolish their responses to low-glucose stimulation (Extended Data Fig. 6e-h), though it modestly suppressed cortical NE release (Extended Data Fig. 6i, j) and partially affected hypoglycemic coma (Extended Data Fig. 6k-m). In the LC NE neurons, no changes in action potential firing or membrane potential level to low-glucose stimulation were observed, as measured by whole-cell recording from GFP+ LC neurons in TH-GFP mice (Extended Data Fig. 6n, o). Collectively, these findings identify AMPK as a principal energy-sensing mechanism through which LC-projecting NTS neurons drive hypoglycemia-induced NE release and oppose coma.

### Brain-wide NE release in CO2-induced hypercapnic coma

Our preceding results indicated that hypoglycemia engages AMPK-dependent activation of LC-projecting NTS neurons, thereby recruiting LC NE neurons and widespread NE release to counteract coma. We next asked whether this neuromodulatory pathway is specific to hypoglycemic coma or represents a broader mechanism for counteracting metabolic coma. To answer this question, a mouse paradigm for inducing and quantifying CO2-induced hypercapnic coma was established (<u>Supplementary Video 2</u>). Mice were placed in a chamber and exposed for 30 min to a gas mixture containing 40% CO2, 20% O2, and 40% N2 at a flow rate of 10 L/min. This exposure induced LORR in mice within approximately 1 min (Fig. 5a), with EEG waveforms and spectral power distributions distinct from those during other brain states (Fig. 5b). The EEG ratio fluctuated as mice transitioned between wakefulness (W) and NREM sleep (N) (Fig. 5c<u>, *left*</u>), but declined rapidly upon CO_2_ exposure, reached a minimum, and then gradually increased after cessation of CO_2_ delivery (Fig. 5c<u>, *left*</u>). Across 15 mice, the minimum EEG ratio during LORR was significantly lower than values recorded in all other brain states (Fig. 5c<u>, *right*</u>).

**Fig. 5.**
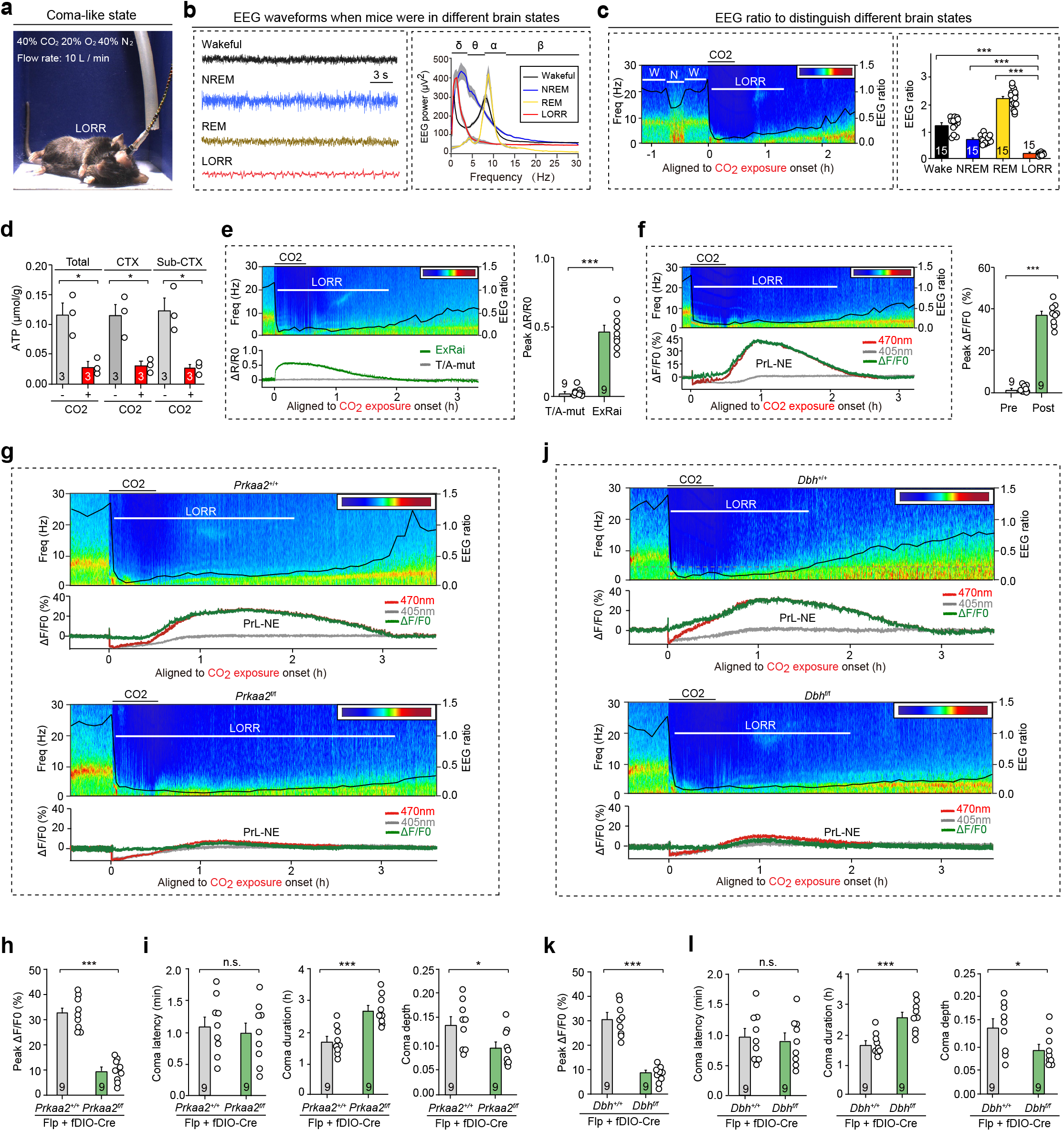
AMPK-dependent NE release is conserved in CO_2_-induced hypercapnic coma. **(a)** Representative image showing LORR in a mouse subjected to CO_2_-induced hypercapnic coma. **(b)** Representative EEG traces (*left*) and EEG power distributions across frequencies during wakefulness, NREM/REM sleep, and LORR (*right*). **(c)** EEG heatmap, EEG ratio curve, and LORR timing in a representative mouse (*left*) and quantification of EEG ratios during wakefulness, NREM/REM sleep, and LORR timing in 15 mice (*right*). W, wakefulness. N, NREM sleep. **(d)** Quantification of brain ATP levels in mice with or without CO_2_-induced hypercapnic coma. **(e)** Representative traces (*left*) and quantification (*right*) of ExRai AMPKAR 470/405 ratio changes, showing an increase in AMPK activity in LC-projecting NTS neurons after CO_2_-induced hypercapnic coma. T/A-mut was used as a negative control. **(f)** Representative traces (*left*) and quantification (*right*) of GRAB-NE2m fluorescence changes, showing an increase in GRAB-NE2m signals in the PrL after CO_2_-induced hypercapnic coma. **(g, h)** Representative traces (g) and quantification (h) of GRAB-NE2m signals during CO_2_-induced hypercapnic coma in mice with or without *Prkaa2* deletion in LC-projecting NTS neurons. Note red and grey curves represent fluorescence changes under 470-nm and 405-nm excitation, respectively. **(i)** Quantification of effects of *Prkaa2* deletion in LC-projecting NTS neurons on CO_2_-induced hypercapnic coma latency (*left*), duration (*middle*), and depth (*right*). **(j, k)** Representative traces (j) and quantification (k) of GRAB-NE2m signals during CO_2_-induced hypercapnic coma in mice with or without *Dbh* deletion in LC NE neurons. **(l)** Quantification of effects of *Dbh* deletion in LC NE neurons on CO_2_-induced hypercapnic coma latency (*left*), duration (*middle*), and depth (*right*). Scale bars are labeled in graphs. Numbers of mice (c-f, h, i, k, l) are indicated in graphs. Data in (c-f, h, i, k, l) are mean ± SEM. Statistical analyses (c-f, h, i, k, l) were performed using Student *t*-test (*** *P* < 0.001; * *P* < 0.05; n.s. *P* > 0.1). For *P*-values, see Supplementary Table 3.

We next examined whether hypercapnia, like hypoglycemia, also induces energy depletion, AMPK activation, and cortical NE release. Notably, CO_2_ exposure for 30 min significantly reduced arterial blood pH from 7.25 ± 0.02 to 6.58 ± 0.03 (*n* = 9 mice). This acidified state was associated with marked ATP reduction across cortical and subcortical brain regions (Fig. 5d), likely reflecting impaired ATP synthesis under severe acidosis. Fiber photometry recordings of AMPKAR in LC-projecting NTS neurons revealed a rapid increase in the 470/405 ratio after CO_2_ exposure (Fig. 5e). In parallel, GRAB-NE2m recordings in the PrL revealed a gradual and robust increase in GRAB-NE2m signals after CO_2_ exposure (Fig. 5f). Because the hypercapnia-induced pH changes can affect fluorescence measurements, we recorded fluorescence signals evoked by excitation at 470 nm (*red*) and 405 nm (*gray*), calculated the normalized fluorescence changes (ΔF/F0, *green*), and thereby corrected experimental artifacts caused by pH changes (Extended Data Fig. 7a)^27^. These findings indicate that hypercapnia, like hypoglycemia, is accompanied by brain energy depletion, AMPK activation in LC-projecting NTS neurons, and cortical NE release.

The requirement for this pathway during hypercapnic coma was then tested. In mice subjected to CO_2_-induced hypercapnic coma, *Prkaa2* deletion in LC-projecting NTS neurons significantly suppressed cortical NE release (Fig. 5g, h). This manipulation did not alter coma latency, but markedly prolonged coma duration and increased coma depth (Fig. 5i). Furthermore, deletion of *Dbh* in LC NE neurons significantly reduced cortical NE release (Fig. 5j, k). Although *Dbh* deletion did not affect coma latency, it prolonged coma duration and increased coma depth (Fig. 5l). Together, these results show that hypercapnia, similar to hypoglycemia, recruits an AMPK-dependent NTS-LC pathway that promotes cortical NE release and limits coma duration and depth.

### VNS recruits widespread NE release

VNS has shown promise for improving arousal in patients with disorders of consciousness^13,14^, but the underlying mechanisms remain incompletely defined. We therefore investigated whether widespread NE release contributes to VNS-evoked arousal from coma in mice. First, a viral tracing strategy was employed to delineate the neural circuit linking vagal afferents to LC NE neurons. In Ai14 mice, AAV1-Cre and AAV-hSyn-DIO-EYFP were injected into the nodose ganglion (NG) and NTS, respectively (Fig. 6a). At 14 days post-injection, abundant tdTomato+ neurons were detected in the NG, confirming efficient AAV1-Cre transduction (Fig. 6b<u>, *left*</u>). The NTS neurons that received monosynaptic innervation of vagal afferents were labeled by EYFP (Fig. 6b<u>, *right*</u>). EYFP+ axons from these NTS neurons were observed in the LC (Fig. 6c). These anatomical data indicate that LC-projecting NTS neurons from a relay between vagal afferents and LC NE neurons.

**Fig. 6.**
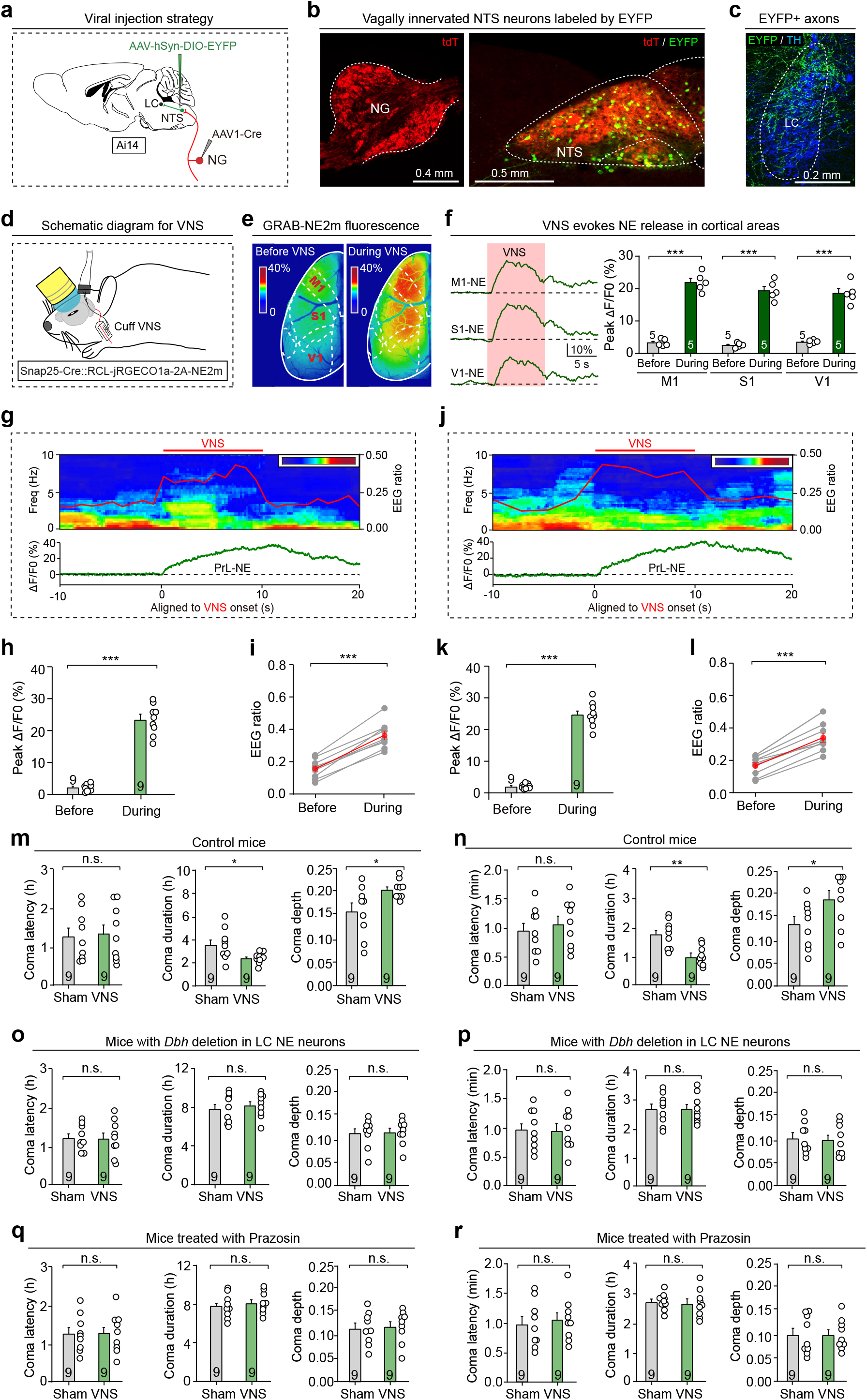
VNS promotes coma recovery by recruiting brain-wide NE release. **(a)** Schematic showing viral injection strategy used to label vagally innervated NTS neurons with EYFP. **(b)** Representative micrographs showing tdTomato-labeled neurons in the NG (*left*) and EYFP-labeled vagally innervated neurons in the NTS (*right*). **(c)** Representative micrograph showing EYFP+ axons intermingled with TH+ LC NE neurons. **(d)** Schematic showing VNS with a cuff electrode in Snap25-Cre::RCL-jRGECO1a-2A-NE2m mice. **(e)** Representative images showing mesoscopic imaging of GRAB-NE2m signals in cortical areas of Snap25-Cre::RCL-jRGECO1a-2A-NE2m mice before and during VNS. **(f)** Representative traces (*left*) and quantification of peak ΔF/F0 (*right*) of GRAB-NE2m signals in M1, S1, and V1. **(g, j)** Representative traces of EEG ratio and GRAB-NE2m signals in the PrL before, during, and after VNS in mice with insulin-induced hypoglycemic coma (g) and CO_2_-induced hypercapnic coma (j). **(h, k)** Quantification of peak ΔF/F0 of GRAB-NE2m signals in the PrL before and during VNS in mice with insulin-induced hypoglycemic coma (h) or CO_2_-induced hypercapnic coma (k). **(i, l)** Quantification of EEG ratio before and during VNS in mice with insulin-induced hypoglycemic coma (i) or CO_2_-induced hypercapnic coma (l). **(m, n)** Quantification showing effect of VNS on insulin-induced hypoglycemic coma (m) or CO_2_-induced hypercapnic coma (n) latency (*left*), duration (*middle*), and depth (*right*) in wild-type mice. **(o, p)** Quantification showing effect of VNS on insulin-induced hypoglycemic coma (o) or CO_2_-induced hypercapnic coma (p) latency (*left*), duration (*middle*), and depth (*right*) in mice with *Dbh* deletion in LC NE neurons. **(q, r)** Quantification showing effect of VNS on insulin-induced hypoglycemic coma (q) or CO_2_-induced hypercapnic coma (r) latency (*left*), duration (*middle*), and depth (*right*) in mice pretreated with α1 receptor antagonist Prazosin. Scale bars are labeled in graphs. Numbers of mice (f, h, i, k, l, m-r) are indicated in graphs. Data in (f, h, i, k, l, m-r) are mean ± SEM. Statistical analyses (f, h, i, k, l, m-r) were performed using Student *t*-test (*** *P* < 0.001; ** *P* < 0.01; * *P* < 0.05; n.s. *P* > 0.1). For *P*-values, see Supplementary Table 3.

We then implanted a cuff electrode on the right vagus nerve to assess the effect of VNS during hypoglycemic and hypercapnic coma. In Snap25-Cre::RCL-jRGECO1a-2A-NE2m mice, mesoscopic imaging was used to monitor cortical GRAB-NE2m responses to VNS (Fig. 6d, e). Brief VNS (20 Hz, 1 mA, 10 s) evoked NE release across multiple cortical regions, as evidenced by marked increases in GRAB-NE2m signals (Fig. 6f). Simultaneous EEG recordings and fiber recording of PrL GRAB-NE2m signals further showed that VNS induced robust PrL NE release, accompanied by a significant increase in the EEG ratio during both hypoglycemic (Fig. 6g-i) and hypercapnic coma (Fig. 6j-l). However, VNS did not significantly alter blood glucose level in hypoglycemic mice (Extended Data Fig. 7b, c), indicating that this stimulation modulates cortical NE release and brain state without measurably changing systemic glucose availability. When mice entered hypoglycemic coma, repeated VNS (20 Hz, 1 mA, 10 s) once per minute for 2 h significantly shortened coma duration and reduced coma depth (Extended Data Fig. 7d <u>and</u> Fig. 6m). Repeated VNS for 0.5 h produced similar effects in hypercapnic coma, shortening coma duration and attenuating coma depth (Extended Data Fig. 7e <u>and</u> Fig. 6n).

To determine whether the anti-coma effect of VNS requires NE release, we deleted *Dbh* selectively in LC NE neurons. After *Dbh* deletion in LC NE neurons (Fig. 2l, m), VNS failed to alter coma duration or depth in mice with hypoglycemic coma (Extended Data Fig. 7f <u>and</u> Fig. 6o) or hypercapnic coma (Extended Data Fig. 7g <u>and</u> Fig. 6p). These results suggest that NE release from LC NE neurons is required for VNS-modulated coma recovery. We next examined which adrenergic receptor mediates the effect of VNS-induced NE release. In control experiments, mice received intraperitoneal injections of adrenergic receptor agonists. Activation of α1-adrenergic receptors with phenylephrine significantly shortened insulin-induced coma duration, whereas agonists targeting other adrenergic receptors had minimal effects (Extended Data Fig. 7h). Conversely, pretreatment with prazosin (6 mg/kg), an α1-adrenergic receptor antagonist, significantly prolonged coma duration (Extended Data Fig. 7i). In hypoglycemic comatose mice pretreated with prazosin, VNS failed to alter coma duration or coma depth (Extended Data Fig. 7j <u>and</u> Fig. 6q). Similarly, α1 antagonist pretreatment blocked the anti-coma effect of VNS in hypercapnic comatose mice (Extended Data Fig. 7k <u>and</u> Fig. 6r). These results suggest that VNS recruits LC-derived NE release to promote coma recovery through α1-adrenergic receptor signaling.

Together, these findings define an endogenous neuromodulatory mechanism that counteracts metabolic coma in mice. Strong metabolic stress, including insulin-induced hypoglycemia and CO_2_-induced hypercapnia, drives mice into coma-like states (Fig. 7a). Hypoglycemia and hypercapnia impair ATP synthesis in NTS neurons, leading to an increase in AMPK activity and subsequent activation of LC-projecting NTS neurons (Fig. 7b<u>, *left*</u>). These neurons then excite LC NE neurons through a glutamatergic NTS-LC circuit, triggering widespread NE release across the brain and elevating arousal to oppose coma (Fig. 7b<u>, *right*</u>). Notably, VNS can recruit this endogenous neuromodulatory pathway, thereby facilitating recovery from metabolic coma (Fig. 7b<u>, *right*</u>).

**Fig. 7.**
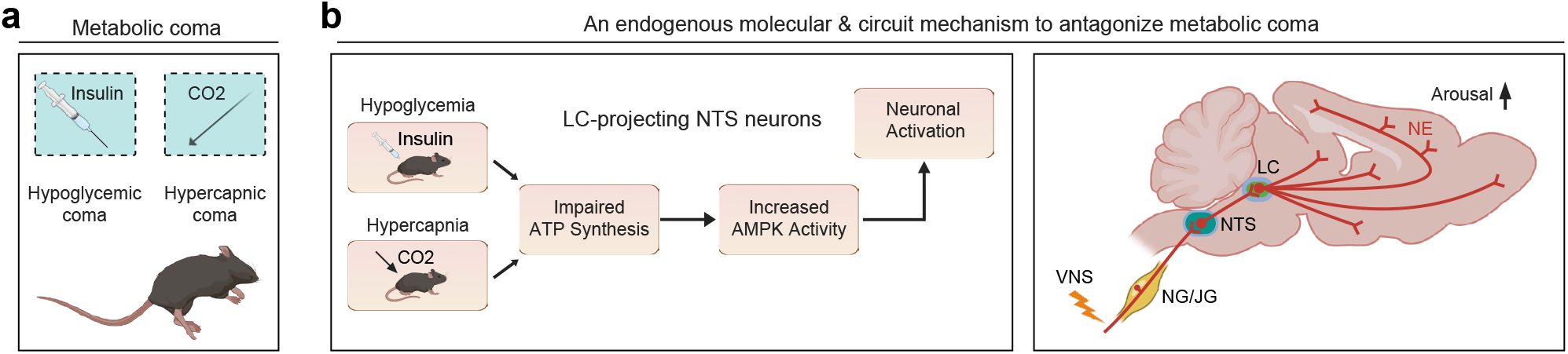
Summary graph of this study. **(a)** Summary of experimental paradigms for metabolic coma induction in mice. **(b)** Schematic summary showing that hypoglycemia and hypercapnia induce AMPK-dependent activation of LC-projecting NTS neurons (*left*), leading to widespread NE release across the brain to counteract coma (*right*). VNS may recruit this anti-coma mechanism to promote post-coma arousal (*right*).

## Discussion

This study identified an energy-sensitive noradrenergic mechanism that limits the duration and depth of metabolic coma in mice. Hypoglycemia and hypercapnia increased AMPK activity in LC-projecting NTS neurons, activated LC NE neurons, and triggered widespread NE release across the brain. Disruption of this pathway prolonged coma and increased coma depth, whereas VNS engaged the same circuitry and promoted recovery. These findings suggest that metabolic stress recruits a noradrenergic arousal mechanism that supports recovery from coma.

### Paradigms to induce and quantify metabolic coma in mice

Although coma is a well-known pathological brain state defined by loss of consciousness^24^, the molecular, cellular, and circuit processes that enable recovery remain poorly resolved. A major barrier has been the lack of experimentally tractable mouse paradigms that permit controlled induction, quantitative assessment, and mechanistic dissection of coma. In this study, we established two mouse models of metabolic coma: insulin-induced hypoglycemic coma (Fig. 1) and CO_2_-induced hypercapnic coma (Fig. 5). In both models, LORR, a classical behavioral index of arousal state in rodents^34^, was used to quantify coma latency and duration. EEG recordings provided an additional physiological measure, with the θ/δ power ratio used to index coma depth. The integration of latency, duration, and depth establishes a quantitative framework for defining coma-like states in mice and for comparing mechanistic interventions across distinct metabolic insults.

Insulin-induced hypoglycemic coma shares several physiological features with torpor and hibernation, two hypometabolic states elicited by energy deficiency and marked by sustained reductions in body temperature^35,36^. However, these states differ fundamentally in reversibility and arousability. Animals in torpor or hibernation can be externally aroused, whereas humans and animals in hypoglycemic coma cannot be readily awakened^37^. Thus, insulin-induced hypoglycemic coma may engage selected energy-conservation pathways that overlap with torpor and hibernation, but the mechanisms governing entry, maintenance, and recovery are not equivalent across these states.

### Neuronal sensing of energy deficiency

This study, together with previous work^30,38,39^, indicates that the NTS contains discrete neuronal populations capable of sensing cellular energy deficiency. These glucose-sensing neurons appear to organize distinct downstream circuits that couple metabolic state to specific physiological outputs. For example, the NTS–DMV pathway regulates blood glucose homeostasis^30^, whereas the TH⁺ NTS–ARC pathway may contribute to feeding regulation^39^. The present findings identify the NTS–LC pathway as an additional energy-sensitive circuit that links metabolic stress to arousal control. Together, these findings suggest that the NTS serves as a central metabolic-sensing hub that detects energy deficiency and distributes this information through specialized downstream circuits to coordinate adaptive physiological responses.

A key question is how NTS neurons detect energy deficiency at the molecular level. Our findings showed that deletion of *Prkaa2* specifically in LC-projecting NTS neurons abolished low-glucose responsiveness. This genetic evidence, together with previous pharmacological studies^30,33^, supports a critical role for AMPK in mediating physiological responses to energy deficiency.

The mechanism through which increased AMPK activity excites LC-projecting NTS neurons remains unresolved. Based on previous work in the hypothalamus^31^, AMPK in NTS neurons may increase neuronal firing under low-glucose conditions through the ion channel Ano4—a hypothesis that warrants future investigation.

### Brain-wide NE release during energy deficiency

LC neurons are established regulators of arousal^40,41^ and sleep^42,43^, likely through NE-dependent modulation of cortical neuronal networks^44,45^. However, the coma-associated NE release identified here differs markedly from conventional state-dependent NE dynamics. GRAB-NE2m signals increased during the wake-to-coma transition, in contrast to the signal decline observed during the wake-to-sleep transition (Extended Data Fig. 3). These coma-associated NE responses also persisted for 2–3 h and reached a magnitude 4- to 10-fold greater than NE fluctuations observed during wake-sleep cycles (Extended Data Fig. 3). Based on its duration, amplitude, and brain-wide distribution, we refer to this response as an “NE storm”. Intriguingly, a similar NE storm also emerged following the onset of hypercapnic coma. Genetic ablation of NE synthesis by *Dbh* deletion in LC NE neurons effectively blocked the NE storm and markedly increased both coma duration and depth, supporting the conclusion that this response functions as an intrinsic anti-coma mechanism.

### VNS facilitates post-coma arousal by recruiting the “NE storm”

VNS induces widespread cortical excitation via multiple arousal-promoting pathways^46^. For example, VNS evokes phasic firing of LC NE neurons and facilitates central NE release^47^, while also activating cholinergic neurons in the basal forebrain^48^. The present findings showed that VNS promotes arousal from coma by inducing brain-wide NE release through an α1-adrenergic receptor-dependent mechanism. However, these data do not exclude potential contributions from the cholinergic system to VNS-modulated coma recovery. We also do not exclude the contributions of vagal efferents to VNS-modulated coma recovery.

Whether VNS also regulates blood glucose level is another important consideration. Under the stimulation parameters used here, VNS did not elevate blood glucose levels in hypoglycemic comatose mice, suggesting that its arousal-promoting effects on hypoglycemic coma are not mediated by counterregulatory enhancement of systemic glucose availability.

## Methods

### Mice

All mouse experiments were performed in accordance with protocols approved by the Administrative Panel on Laboratory Animal Care (NIBS2026M0001) at the National Institute of Biological Sciences (NIBS), Beijing, China. C57BL/6J mice were used together with the following genetically modified lines: Snap25-Cre^49^, Dbh-Cre^50^, Slc17a7-Cre^49^, Slc32a1-Cre^51^, Th-Flp^52^, Ai14^53^, RCL-jRGECO1a-2A-NE2m^27^, and TH-GFP^54^. These lines were obtained from the Jackson Laboratory (JAX Mice and Services). Mice were maintained on a circadian 12-h light/12-h dark cycle with food and water available *ad libitum*. Before viral injection, mice were group-housed (3–5 animals/cage) and then separated 3 days before surgery. After viral injection, each mouse was housed individually for at least 3 weeks before subsequent experiments. To minimize potential sex-specific differences, only male mice were used. Additional information on mouse lines is provided in the Supplementary Table 1.

### Viral vectors and stereotaxic injections

#### Viral vectors

Viral particles were purchased from Shanghai Taitool Bioscience Inc. and BrainCase Inc. Adeno-associated virus (AAV) serotype 2/9 (AAV2/9) was used for AAV-mediated delivery. Before dilution, AAV titers ranged from 0.8 to 1.5 × 10^13^ viral particles/mL, and the final titer used for injection was 5 × 10^12^ viral particles/mL. The rabies virus (RV) titer was 5.0 × 10^8^ viral particles/mL, and the pseudorabies virus (PRV) titer was 2 × 10^9^ PFU/mL. All AAV, RV, and PRV vectors used in this study are listed in Supplementary Table 1.

#### Brain stereotaxic injections

Stereotaxic viral injections into the brain were performed as described previously^55^. Mice were anesthetized by intraperitoneal injection of tribromoethanol (250 mg/kg), and the brain surface above the LC, PrL, M1, LH, PAG, NTS and S1 were exposed using standard surgical procedures. The coordinates for LC injection were: bregma −5.40 mm, lateral ±0.75 mm, and dura −2.50 mm. The coordinates used for PrL injection were: bregma +2.80 mm, lateral ±0.25 mm, and dura −0.75 mm. The coordinates used for M1 injection were: bregma +1.98 mm, lateral ±1.50 mm, and dura −0.75 mm. The coordinates used for LH injection were: bregma -1.58 mm, lateral ±1.10 mm, and dura −4.75 mm. The coordinates used for PAG injection were: bregma -4.60 mm, lateral ±0.35 mm, and dura −1.75 mm. The coordinates used for NTS injection were: bregma -7.56 mm, lateral ±0.50 mm, and dura −2.50 mm. The coordinates used for S1 injection were: bregma -0.46 mm, lateral ±2.0 mm, and dura −0.50 mm. Viral vectors were delivered through a glass pipette connected to a Nanoliter Injector 201 (World Precision Instruments, Inc.) at a slow flow rate of 0.15 μL/min to minimize local brain tissue damage. The pipette was left in place for at least 20 min before withdrawal. After injection, the skin was sutured, and antibiotics were applied to the surgical wound. Bilateral injections were performed for chemogenetic activation or inactivation and for anterograde or retrograde tracing. Unilateral injections were performed for optogenetic activation.

#### AAV injection in the NG

AAV injection into the nodose ganglion (NG) was performed following previously published protocols^56^. In brief, anesthetized mice were placed in ventral recumbency, and an approximate 1.5-cm incision was made along the ventral neck surface to expose the NG. Pulled glass pipettes with a tip diameter of around 15 μm were filled with AAV, inserted into the target area, and used to deliver virus at 500 nL/min with a Nanoliter Injector 201 (World Precision Instruments, Inc.). The pipette was left in place for at least 20 min before withdrawal. After injection, the skin was sutured, and antibiotics were applied to the surgical wound.

### Optical fiber implantation

Optical fiber implantation was performed as described previously^55^. At 30 min post-AAV injection, a ceramic ferrule containing an optical fiber (200 µm in diameter, NA 0.37) was implanted, with the fiber tip positioned above the LC (bregma −5.40 mm, lateral ±0.75 mm, dura −2.50 mm), PrL (bregma +2.80 mm, lateral ±0.25 mm, dura −0.75 mm), M1 (bregma +1.98 mm, lateral ±1.50 mm, dura −0.75 mm), LH (bregma -1.58 mm, lateral ±1.10 mm, dura −4.75 mm), PAG (bregma -4.60 mm, lateral ±0.35 mm, dura −1.75 mm) or NTS (bregma −7.56 mm, lateral ±0.50 mm, dura −2.50 mm). The ferrule was secured to the skull with dental cement. After implantation, the skin was sutured, and antibiotics were applied to the wound. Fiber photometry recordings and optogenetic activation experiments were conducted 3 weeks after implantation.

### Preparation of behavioral measurements

After AAV injection and optical fiber implantation, mice were housed individually for 3 weeks. Before behavioral testing, mice were handled daily by the experimenters for at least 3 days. On the day of testing, mice were transferred to the behavioral room and habituated for 3 h before the experiments began. All apparatuses were cleaned with 20% ethanol to eliminate odor cues from other mice. Behavioral tests were conducted during the same circadian period (13:00–19:00). All behaviors were scored by investigators blinded to experimental treatments.

### Induction of metabolic coma and LORR testing in mice

#### Insulin-induced hypoglycemic coma in mice

Mice were fasted for 24 h and placed in an arena (7.5 cm × 7.5 cm × 15 cm) for continuous EEG recording. Insulin (12 U/kg; NovoRapid, China) was diluted 1:100 in saline and administered by intraperitoneal injection. Loss of righting reflex (LORR) was assessed by rotating the arena 180° with a motorized device. Given the relatively long latency of insulin-induced coma, LORR was tested every 10 min. Mice were considered to have lost the righting reflex when all four paws failed to contact the ground after arena rotation. Testing was terminated once LORR was observed, and the experiment ended after spontaneous recovery of the righting reflex. Latency to hypoglycemic coma was defined as the time from insulin injection to LORR. Duration of hypoglycemic coma was defined as the time from LORR to recovery of righting reflex.

#### LORR testing in insulin-induced coma

Following intraperitoneal injection of insulin (12 U/kg), mice were placed in a freely rotatable arena (7.5 cm × 7.5 cm × 15 cm) and continuously video-recorded. During the experiment, the arena was rotated 180° every 10 min to assess the righting reflex. Mice were considered to have developed LORR when all four paws failed to contact the ground after rotation. Once LORR was observed, reflex testing was ceased, and video recording continued until spontaneous recovery of the righting reflex.

#### CO_2_-induced hypercapnic coma in mice

Mice were placed in an arena (7.5 cm × 7.5 cm × 15 cm) for continuous EEG recording. The arena was sealed except for inlet and outlet ventilation ports. Mice were exposed for 30 min to 40% CO**_2_** gas (40% CO**_2_**, 20% O**_2_**, 40% N**_2_**) at a flow rate of 10 L/min, after which gas delivery was terminated. LORR was assessed by rotating the arena 180° with a motorized device. Given the relatively short latency of CO**_2_**-induced coma, LORR was tested every 30 s. Mice were considered to have lost the righting reflex when all four paws failed to contact the ground following arena rotation. Testing was halted once LORR was observed, and the experiment ended after spontaneous recovery of the righting reflex. Latency to hypercapnic coma was defined as the time from the onset of CO**_2_** exposure to LORR. Duration of hypercapnic coma was defined as the time from LORR to recovery of righting reflex.

#### LORR testing in CO_2_-induced coma

Mice were placed in a sealed, freely rotatable arena (7.5 cm × 7.5 cm × 15 cm) equipped only with inlet and outlet ventilation ports. The arena was infused with 40% CO_2_ for 30 min. Given the short latency of CO_2_-induced coma, the arena was rotated 180° every 30 s to assess the righting reflex. Mice were considered to have developed LORR when all four paws failed to contact the ground following rotation. Once LORR was observed, reflex testing was stopped, and recording continued until spontaneous recovery of the righting reflex.

### EEG recording and analyses

#### EEG electrode implantation

Mice were anesthetized with tribromoethanol (250 mg/kg) via intraperitoneal injection and secured in a stereotaxic apparatus. After removal of the scalp and periosteum, screw electrodes were implanted at the following coordinates relative to bregma: frontal association cortex (FrA, recording electrode): AP +2.58 mm, ML +1.5 mm; right hippocampus (reference electrode): AP −2.0 mm, ML −1.37 mm; and the avascular midline region of the cerebellum (ground electrode): AP −6.3 mm, ML 0 mm. The electrodes were fixed with dental cement. The EEG connector pedestal was soldered to the screw wires and further secured with dental cement. Mice were allowed to recover for 7 days before EEG recording.

#### EEG signal recording

EEG signals were acquired using LabChart software (ADInstruments, USA) at a sampling rate of 400 Hz. During acquisition, a 0.3–100 Hz band-pass filter was applied, and a notch filter was enabled to suppress power frequency interference. The input range was set to ±2 mV. Behavioral states, including wakefulness, NREM sleep, REM sleep, and coma, were determined based on standard EEG characteristics combined with behavioral observations.

#### Power spectral density (PSD) analysis

For each behavioral state, artifact-free 30-s EEG segments were selected to ensure the data were free of motion artifacts or signal distortion. Power spectral density was calculated using fast Fourier transform (FFT) with a 1 024-point FFT length, Hamming window, 93.75% window overlap, and removal of the DC component. The analysis frequency range was 0–30 Hz, with a frequency resolution of approximately 0.39 Hz, calculated from the sampling rate divided by the FFT length. When required, power within each frequency band was normalized to total power across the 0–30 Hz range.

#### Analysis of EEG ratio

To compare EEG patterns across conscious states, the θ/δ power ratio was calculated as the EEG ratio. For wakefulness, NREM sleep, REM sleep, and LORR, artifact-free 30-s EEG segments were selected, and θ-band power (4–8 Hz) and δ-band power (0.5–4 Hz) were calculated. The EEG ratio was defined as θ power divided by δ power. Across the entire coma induction experiment, the EEG ratio was calculated in sequential segments at 5-min intervals, and the resulting time-course curve was used to characterize dynamic transitions in conscious state.

#### EEG time-frequency heatmap

To assess time-dependent changes in EEG spectral structure, time-frequency heatmaps were generated using a custom MATLAB script. Raw LabChart data were exported in .wav format and analyzed by short-time Fourier transform (STFT) using parameters consistent with those applied in PSD analysis, including a sampling rate of 400 Hz and a frequency range of 0–30 Hz. Power was calculated for each time-frequency unit and visualized as a heatmap, with time on the horizontal axis, frequency (0–30 Hz) on the vertical axis, and color indicating power intensity. To enhance visualization contrast, power values were log-transformed and expressed in decibels (dB). Heatmaps were used to visualize the temporal evolution of EEG spectral activity across wakefulness, sleep, and coma states.

### Measurements of hypoglycemia-related symptoms

#### Blood glucose level

Mice implanted with EEG electrodes were also implanted with a DSI glucose telemeter (HD-XG) and allowed to recover for 3 days before experiments. After connection to the EEG recording system, mice were placed in an arena (7.5 cm × 7.5 cm × 15 cm) positioned directly above the receiver. Blood glucose dynamics were recorded using Ponemah software and calibrated with Roche blood glucose test strips.

#### Heart rate and breathing rate

Mice implanted with EEG electrodes were also implanted with a DSI heart rate telemeter (HD-X11) and allowed to recover for 3 days before experiments. After connection to the EEG recording system, mice were placed in a whole-body plethysmography (WBP) chamber (DSI, 601-1425-001) positioned directly above the receiver. EEG, respiration, and heart rate changes were simultaneously recorded using LabChart, FinePointe, and New Ponemah Acquisitions software, respectively.

#### Body temperature

After connection to the EEG recording system, mice were placed in an arena (7.5 cm × 7.5 cm × 15 cm), and body temperature was monitored in real time using a thermal imaging system (Fotric, China).

#### Measurement of pupil diameter changes

Mice were anesthetized with tribromoethanol (250 mg/kg) via intraperitoneal injection and secured in a stereotaxic apparatus. A headplate was implanted, and mice were allowed to recover for 1 day before testing. During testing, mice were head-fixed on a roller and imaged with an infrared camera. Changes in pupil diameter before and after light stimulation (50 mW, 5 s) were recorded in awake and insulin-induced coma states using ISCAN.

### Measurement of brain glucose levels

Intracerebral glucose level was measured according to published protocols^57^. After 24 h of fasting, mice were anesthetized with tribromoethanol (250 mg/kg) via intraperitoneal injection and secured in a stereotaxic apparatus. A working electrode (MT10, China) was implanted into the prelimbic/infralimbic cortex (PrL/IL) brain region (AP +1.8 mm, ML +0.3 mm, DV −2.5 mm), and a reference electrode and an auxiliary electrode (MT51, China) were implanted nearby; EEG electrodes were implanted during the same procedure, and all electrodes were fixed with dental cement. Intracerebral glucose levels were measured electrochemically using a neurochemical analysis system (MT200, China). This system records current changes that can be converted into glucose concentrations. The recording voltage was set to 0.13 V, with a sampling interval of 0.1 s. After recovery from anesthesia, simultaneous recording of EEG activity and intracerebral glucose levels was initiated.

### Measurement of brain ATP levels

#### Measurement of ATP levels after insulin injection

Wild-type mice were fasted for 24 h and then received an intraperitoneal injection of insulin (12 U/kg). One hour post-injection, mice were anesthetized with tribromoethanol (250 mg/kg) via intraperitoneal injection, and brains were rapidly harvested on ice. Cortical and subcortical tissues were dissected separately. Equal amounts of tissue from control and insulin-treated mice were processed, and ATP levels were measured according to the protocol for the Enhanced ATP Assay Kit (Beyotime S0027, China).

#### Measurement of ATP levels after CO_2_-induced hypercapnia

Wild-type mice were exposed to 40% CO_2_ for 30 min. Mice were then anesthetized with tribromoethanol (250 mg/kg) via intraperitoneal injection, and brains were rapidly harvested on ice. Cortical and subcortical tissues were dissected separately. Equal amounts of tissue from control and CO_2_-exposed mice were processed, and ATP levels were measured according to the protocol for the Enhanced ATP Assay Kit (Beyotime S0027, China).

#### Measurement of ATP levels using enhanced ATP assay kit

Tissue samples were lysed by adding lysis buffer at approximately 100–200 μL per 20 mg tissue, followed by thorough homogenization with a glass homogenizer. Sufficient homogenization was performed to ensure complete lysis of the tissue. Lysates were centrifuged at 12 000 ×*g* for 5 min at 4°C and the supernatant was collected for subsequent ATP measurement. ATP standards were prepared by diluting the ATP standard solution in ATP assay lysis buffer to generate a concentration gradient for the standard curve. ATP assay working solution was prepared immediately before measurement, with 100 μL required for each sample or standard. ATP assay reagent was diluted with ATP assay reagent diluent at a ratio of 1:4. For each measurement, 100 μL of ATP assay working solution was added to each detection well or tube and incubated at room temperature for 3–5 min to reduce background ATP. Subsequently, 20 μL of sample or standard was added and mixed with a micropipette, with luminescence measured after an interval of at least 2 s using a multi-mode microplate reader (Spark, TECAN). ATP concentrations were calculated from the standard curve.

### Measurement of blood pH in CO₂-Induced hypercapnia

Wild-type mice were ventilated to gas mixture (40% CO₂, 20% O₂, 40% N₂) for 30 minutes to induce hypercapnia. At the end of the ventilation period, arterial blood samples were rapidly withdrawn from the left cardiac ventricle using a heparinized syringe and immediately analyzed with a portable blood gas analyzer (i-STAT, Abbott, USA). Blood gas parameters, with a particular focus on pH, were recorded to assess the systemic physiological responses to the hypercapnic challenge.

### Two-photon imaging in mouse cortex

#### Two-photon imaging during tail-pinch stimulation in S1 before and after coma

Methods for two-photon imaging followed our previous study^58^. AAV-hSyn-jGCaMP7f was injected into the S1 region of wild-type mice. At 21 days post-injection, mice were anesthetized with tribromoethanol (250 mg/kg) via intraperitoneal injection and secured in a stereotaxic apparatus. After careful removal of the scalp and overlying skull muscles, a custom-made chamber was adhered to the skull with cyanoacrylate and fixed with dental cement. A square cranial window (0.25 cm × 0.25 cm) was drilled above S1, the dura mater was carefully removed with fine forceps, and a square coverslip (0.25 cm × 0.25 cm) was placed over the cranial window and fixed with dental cement. Five days later, mice were fasted for 24 h before two-photon imaging. Imaging was performed on a Nikon A1+ MP microscope equipped with a 25× water-immersion objective, using 920-nm excitation and 500–550-nm emission collection. Imaging depth was 200 μm, whereas acquisition frequency was 7.7 Hz. During wakefulness, a 2-min baseline recording was obtained, followed by tail-pinch stimulation with an alligator clip (250 g force) and an additional 2-min recording. Mice then received an intraperitoneal injection of insulin (12 U/kg). After LORR was observed, a 2-min baseline was recorded, the tail was pinched with an alligator clip (250 g force), and imaging continued for another 2 min.

#### Two-photon imaging of GRAB-NE2m in single S1 neurons

AAV-DIO-GRAB-NE2m and AAV-DIO-tdTomato were injected into the S1 cortex of Slc17a7-Cre or Slc32a1-Cre mice. After 21 days, mice were anesthetized with tribromoethanol (250 mg/kg) via intraperitoneal injection and fixed on a stereotaxic apparatus. After careful removal of the scalp and overlying skull muscles, a custom-made chamber was adhered to the skull with cyanoacrylate and fixed with dental cement. A square cranial window (0.25 cm × 0.25 cm) was drilled in the skull above S1, the dura mater was carefully removed with fine forceps, and a square coverslip (0.25 cm × 0.25 cm) was placed over the cranial window and fixed with dental cement. After 5 days, mice were fasted for 24 h before two-photon imaging. Imaging was performed on a Nikon A1+ MP microscope equipped with a 25× water-immersion objective, with 920-nm excitation and 500–550-nm emission collection for GRAB-NE2m, 570-620 nm for tdTomato.

The imaging depth was 200 μm, whereas acquisition frequency was 7.7 Hz. Somatic GRAB-NE2m signals from Slc17a7+ or Slc32a1+ neurons in S1 were recorded before insulin injection and again when the EEG ratio reached its minimum during insulin-induced coma.

#### Analysis and processing of imaging data

Imaging data were analyzed using ImageJ and custom-made MATLAB code. Image stacks were opened in ImageJ and motion-corrected using the Image Stabilizer plugin. Images were overlaid, cells with detectable fluorescence signals were manually outlined, and region-of-interest (ROI) information was saved. ROI files generated in ImageJ were imported into MATLAB and analyzed using the Cell ROI program to extract normalized fluorescence traces for all cells over time. Fluorescence changes were calculated as ΔF/F_0_ (%) = (F − F_0_) / F_0_ × 100.

### Mesoscopic imaging of cortical areas

#### Mesoscopic imaging of the cortex

Mesoscopic imaging procedures followed a previous study^27^. Snap25-Cre::RCL-jRGECO1a-2A-NE2m mice were anesthetized with tribromoethanol (250 mg/kg) via intraperitoneal injection and secured in a stereotaxic apparatus. After removal of the scalp and overlying skull muscles, a custom chamber was attached to the skull with cyanoacrylate and fixed with dental cement. The skull above the dorsal cerebral cortex was removed with a skull drill, the dura mater was carefully removed with fine forceps, and a trapezoidal coverslip covering the entire cerebral cortex was placed over the cranial window and fixed with dental cement. After 3 days of recovery, mice were fasted for 24 h before mesoscopic imaging. Imaging was performed using an Olympus MVX10 macroscope with 405/470-nm excitation and 500–550-nm emission collection equipped with a 2× objective lens (0.5 NA, Olympus, MVPLAPO2XC). Cortical GRAB-NE2m signals were recorded continuously before and after insulin injection.

#### Mesoscopic imaging of VNS-evoked GRAB-NE2m signals

Snap25-Cre::RCL-jRGECO1a-2A-NE2m mice were anesthetized with tribromoethanol and secured in a stereotaxic apparatus. After removal of the scalp and overlying skull muscles, a custom chamber was attached to the skull with cyanoacrylate and fixed with dental cement. The skull above the dorsal cerebral cortex was removed with a skull drill, the dura mater was carefully removed with fine forceps, and a trapezoidal coverslip covering the cerebral cortex was placed on the cranial window and fixed with dental cement. Then the vagal cuff electrode was implanted in mouse as described in the cuff electrode implantation protocol. After 7 days of recovery, mice were fasted for 24 h before imaging. Insulin was administered by intraperitoneal injection, and after mice entered a coma-like state, mesoscopic imaging was performed to record cortical GRAB-NE2m signals before and after VNS. Imaging was performed using an Olympus MVX10 macroscope with 405/470-nm excitation and 500–550-nm emission collection.

#### Analysis of imaging data

Imaging data were analyzed using ImageJ and custom-made MATLAB code. Images were opened in ImageJ and motion-corrected with the Image Stabilizer plugin. ROIs were selected according to cortical region boundaries. ROI information generated in ImageJ was imported into MATLAB and analyzed with a custom ROI analysis program to obtain normalized fluorescence traces over time for each cortical region. The 405-nm channel was used to correct for hemodynamic artifacts. The 405-nm signal was linearly fitted to the 470-nm signal using the least-squares method to generate a scaled control channel, calculated as scaled 405 = a × 405 + b. The corrected fluorescence signal was calculated as F_corrected_ = F_470_ –F_405_fitted_. Fluorescence changes were calculated as ΔF/F_0_ (%) = (F − F_0_) / F_0_ × 100.

### Fiber photometry

#### Instrument configuration

Fiber photometry recordings were performed using a dual-color optical fiber recording system (ThinkerTech, China), which allowed fluorescence signal acquisition with simultaneous correction of motion-related artifacts. A 405-nm light-emitting diode (LED) was used as the control channel, and a 470-nm LED was used to excite GCaMP. Excitation light from each LED was collimated and passed through band-pass filters for the signal channel (470/25 nm) and control channel (405/10 nm). The two excitation paths were then combined through a dichroic mirror and objective lens and coupled into a low-autofluorescence optical fiber patch cord (diameter: 200 μm, numerical aperture: 0.37). The patch cord was connected to the implanted optical fiber ferrule through a rotary joint to permit unrestricted movement during recording. Optical power at the fiber tip was adjusted to 10 μW to minimize photobleaching. Emitted fluorescence (505–550 nm) was collected through the same optical fiber, transmitted back to the system, and detected with a high-sensitivity photodetector. The 405-nm signal, which is insensitive to calcium-dependent fluorescence changes, was used as a real-time reference signal to exclude and correct non-specific noise caused by movement, blood flow, photobleaching or pH changes.

#### Experimental animal preparation

For GCaMP fiber photometry recordings from LC-projecting NTS neurons, wild-type mice received unilateral injection of AAV2-retro-Cre into the LC and unilateral injection of AAV-DIO-jGCaMP7f into the NTS, followed by implantation of optical fibers and EEG electrodes. For AMPK activity recordings from LC-projecting NTS neurons, wild-type mice received unilateral injection of AAV2-retro-Cre into the LC and unilateral injection of AAV-DIO-ExRai-AMPKAR into the NTS, followed by implantation of optical fibers and EEG electrodes. For GCaMP recordings from the LC, Dbh-Cre mice received unilateral injection of AAV-DIO-jGCaMP7f virus into the LC, followed by implantation of optical fibers and EEG electrodes. For NE recordings from the PrL, M1, LH, and PAG, wild-type mice received unilateral or bilateral injections of AAV-hSyn-GRAB-NE2m into the corresponding brain regions, followed by implantation of optical fibers and EEG electrodes. For DA recordings from the PrL, wild-type mice received unilateral injection of AAV-hSyn-GRAB-DA2m into the corresponding brain regions, followed by implantation of optical fibers and EEG electrodes.

#### Fiber photometry recordings and signal analysis

Fiber photometry recordings were performed 21 days after optical fiber implantation. During each experiment, mice were connected to both the EEG recording system and the 405/470-nm dual-color fiber photometry system (ThinkerTech, China). Signals were digitized and recorded at 40 Hz and acquired using manufacturer-provided software. A baseline recording was first obtained during wakefulness, after which mice received an intraperitoneal injection of insulin (12 U/kg) or were exposed to 40% CO_2_ for 30 min. Recording continued until recovery of consciousness. Data were analyzed using Spike2 software. Fiber photometry signals were corrected by the 405 nm reference signal. The 405-nm signal was linearly fitted to the 470-nm signal using the least-squares method to generate a scaled control channel, calculated as scaled 405 = a × 405 + b. The corrected fluorescence signal was calculated as F_corrected_ = F_470_ –F_405_fitted_. The relative fluorescence change was then calculated as ΔF/F_0_ (%) = (F_corrected_ –F_0_) / F_0_ × 100, where F_0_ is the mean value of the corrected fluorescence signal during the baseline period. For peak ΔF/F_0_ calculation, mean ΔF/F_0_ during the 30-s pre-stimulus baseline period was used as the reference. Peak ΔF/F_0_ was defined as the difference between the maximum ΔF/F_0_ value during each hypoglycemic or hypercapnic stimulus and the baseline mean. For AMPKAR fiber photometry signals, the 470/405 excitation ratio was calculated at each time point as described previously^32^. The relative change in this ratio was calculated as ΔR/R_0_ = (R – R_0_) / R_0_, where R_0_ is the mean ratio during the baseline period. To determine peak ΔR/R_0_, the mean ΔR/R_0_ during the 30-s pre-stimulus baseline period was used as the reference. Peak ΔR/R_0_ was defined as the difference between the maximum ΔR/R_0_ value during each hypoglycemic or hypercapnic stimulus and the baseline mean.

#### Comparison of NE signals across behavioral contexts

Twenty-one days after optical fiber implantation, EEG and GRAB-NE2m signals were recorded from the same mice across two contexts: insulin-induced coma after intraperitoneal injection of insulin (12 U/kg), and different sleep stages including NREM and REM. Peak changes in NE signals were quantified and compared across these states.

#### Simultaneous measurement of blood glucose and GRAB-NE2m signals

AAV-hSyn-GRAB-NE2m was unilaterally injected into the PrL of wild-type mice, and optical fibers and EEG electrodes were implanted during the same procedure. After 21 days, mice were implanted with a DSI glucose telemeter (HD-XG) and allowed to recover for 3 days before experimentation. After 24 h of fasting, mice were connected to the EEG and dual-color fiber photometry systems and placed in an arena (7.5 cm × 7.5 cm × 15 cm) positioned directly above the DSI receiver. Baseline signals were first recorded during wakefulness, followed by intraperitoneal injection of insulin (12 U/kg) to induce hypoglycemic coma. EEG activity, blood glucose, and GRAB-NE2m signals were continuously recorded until recovery of consciousness. The blood glucose value corresponding to the onset of GRAB-NE2m signal elevation was quantified and defined as the blood glucose threshold for NE signal elevation.

#### Correction of NE signals during hypercapnia

To correct signal artifacts introduced by motion, nonspecific photobleaching, and pH changes during hypercapnic stimulation, an isosbestic control channel was used to normalize GRAB-norepinephrine (GRAB-NE2m) fluorescence signals^27^. GRAB-NE2m fluorescence signals were acquired under 470-nm excitation, whereas the isosbestic control signal was acquired under 405-nm excitation. The two LED light sources were coupled into a shared optical path via a fiber optic patch cord. Demodulated signals were sampled at 40 Hz and smoothed using a low-pass filter with a 1 s time constant. The 405-nm signal was then linearly fitted to the 470-nm signal using the least-squares method to generate a scaled control channel, calculated as scaled 405 = a × 405 + b. The corrected fluorescence signal was calculated as F_corrected_ = F_470_ –F_405_fitted_. The relative fluorescence change was then calculated as ΔF/F_0_ (%) = (F_corrected_ –F_0_) / F_0_ × 100, where F_0_ is the mean value of the corrected fluorescence signal during the baseline period. For peak ΔF/F_0_ calculation, mean ΔF/F_0_ during the 30-s pre-stimulus baseline period was used as the reference. Peak ΔF/F_0_ was defined as the difference between the maximum ΔF/F_0_ value during each hypercapnic stimulus and the baseline mean.

### Cell type-specific RV tracing

A modified RV-based three-virus system was used to map brain-wide monosynaptic inputs to LC NE neurons^28^. All viruses were packaged and supplied by BrainVTA Inc. (Wuhan, China), including AAV2/9-EF1α-DIO-EGFP-2A-TVA (5 × 10^12^ viral particles/mL), AAV2/9-Ef1α-DIO-RV-G (5 × 10^12^ viral particles/mL), and RV-EnvA-ΔG-tdTomato virus (5.0 × 10^8^ viral particles/mL). A 1:1 mixture of AAV2/9-EF1α-DIO-EGFP-2A-TVA and AAV2/9-Ef1α-DIO-RV-G (200 nL) was bilaterally injected into the LC of Dbh-Cre mice. Two weeks after AAV helper injection, RV-EnvA-ΔG-tdTomato (300 nL) was injected into the same LC sites in a biosafety level-2 lab facility. Starter neurons were identified by co-expression of tdTomato and EGFP and were restricted to the LC. One week after RV injection, mice were perfused with saline, followed by 4% paraformaldehyde (PFA) in phosphate-buffered saline (PBS). Brains were post-fixed in 4% PFA for 8 h and then incubated in PBS containing 30% sucrose until they sank. Coronal brain sections (40 μm) were prepared using a cryostat (Leica CM1900). All sections were collected and stained with 4’,6-diamidino-2-phenylindole (DAPI). Sections were imaged using an Olympus VS120 epifluorescence microscope (10× objective lens) and analyzed with ImageJ. Regional boundaries were defined according to the Allen Brain Atlas, and the fractional distribution of total RV-labeled cells was quantified across subregions.

### Immunohistochemical and RNAscope analyses

#### Immunohistochemical analyses

Mice were anesthetized with isoflurane and sequentially perfused with saline and 4% PFA in PBS. Brains were incubated in PBS containing 30% sucrose until they sank. Cryostat sections (40 μm) were collected, incubated overnight in blocking solution containing PBS, 10% goat serum, and 0.7% Triton X-100, and then treated with primary antibodies diluted in blocking solution for 3–4 h at room temperature. Primary antibodies used for immunohistochemical analysis are shown in Supplementary Table 1. Sections were washed three times with washing buffer containing PBS and 0.7% Triton X-100 before incubation with Cy2-, Cy3-, or Cy5-conjugated secondary antibodies (1:500; Life Technologies Inc.) for 1 h at room temperature. Sections were then washed three times with washing buffer, stained with DAPI, washed with PBS, transferred onto Super Frost slides, and mounted under glass coverslips with mounting media. The sections were imaged using an Olympus VS120 epifluorescence microscope (10× objective lens) or a Zeiss LSM 800 confocal microscope (20× and 60× oil-immersion objective lens). Samples were excited with 488-, 543-, or 633-nm lasers in sequential acquisition mode to avoid signal leakage. Pixel saturation was avoided by monitoring signal intensity under Hi-Lo mode. Confocal images were analyzed with ImageJ.

#### Mapping Fos expression in brain areas

After 24 h of fasting, mice received an intraperitoneal injection of insulin (12 U/kg) to induce coma. After coma and recovery of consciousness, mice were sequentially perfused with saline and 4% PFA. Control mice received an intraperitoneal injection of saline and were perfused after placement in the arena for the same duration. After perfusion, brains were post-fixed in 4% PFA for 8 h, dehydrated in PBS containing 30% sucrose, and sectioned. Sections were blocked with 0.7% PBST containing Triton X-100 and 10% fetal bovine serum (FBS) at room temperature for 2 h, incubated with primary antibody (rabbit c-Fos, 1:1 000) at 4°C overnight followed by incubation at room temperature for 2 h, washed three times with 0.7% PBST, incubated with secondary antibody (goat anti-rabbit 488, 1:500) at room temperature for 2 h, washed three times with 0.7% PBST, stained with DAPI, washed with PBS, and then mounted. Imaging was performed using an Olympus VS120 microscope with a 10× objective lens, and Fos-positive cells per unit area were quantified in different brain regions using ImageJ.

#### RNAscope sample preparation

To verify Gal knockout in the LC, TH-2A-Flp::*Gal*^+/+^ and TH-2A-Flp::*Gal*^f/f^ mice were used. AAV-fDIO-EGFP-2A-Cre was bilaterally injected into the LC brain region, and RNAscope analysis was performed 21 days after viral expression. To verify Prkaa1 knockout in LC-projecting NTS neurons, *Prkaa1*^+/+^ and *Prkaa1^f^*^/f^ mice were used. AAV2-retro-Flp virus was bilaterally injected into the LC, and AAV-fDIO-EGFP-2A-Cre was bilaterally injected into the NTS brain region, with RNAscope analysis performed 21 days after viral expression. To verify Prkaa2 knockout in LC-projecting NTS neurons, *Prkaa2*^+/+^ and *Prkaa2^f^*^/f^ mice were used. AAV2-retro-Flp virus was bilaterally injected into the LC, and AAV-fDIO-EGFP-2A-Cre was bilaterally injected into the NTS brain region, with RNAscope analysis performed 21 days after viral expression.

#### RNAscope staining

For RNAscope analysis, mice were sequentially perfused with DEPC-PBS and DEPC-4% PFA. After perfusion, brains were post-fixed in DEPC-4% PFA for 24 h, dehydrated sequentially with 20% and 30% sucrose prepared in DEPC-PBS, and sectioned coronally through the target brain regions (LC/NTS) at 20 μm. Sections were rinsed with DEPC-PBS, mounted on charged glass slides, dried at −20°C for 2 h, and baked at 60°C for 30 min. Slides were then fixed with DEPC-4% PFA at 4°C for 15 min, dehydrated stepwise in ethanol (50%, 70%, and 100% twice, 5 min per step), treated with hydrogen peroxide at room temperature, and subjected to target retrieval at 99°C for 5 min. Slides were incubated with primary antibodies (rabbit GFP, 1:2 000 alone; or rabbit GFP, 1:2 000 and mouse TH, 1:1 000) at 4°C overnight, followed by incubation at room temperature for 2 h. The slides were then post-fixed in DEPC-4% PFA at room temperature for 30 min, incubated with Protease Plus (Advanced Cell Diagnostics) at 40°C for 30 min, rinsed with DEPC-distilled water, and incubated with mouse Gal, Prkaa1, and Prkaa2 RNAscope probes (Advanced Cell Diagnostics) at 40°C for 2 h. Probe signal amplification and channel-specific labeling with Cy3 (TSA Plus Cy3 Reagent, Akoya Biosciences) were performed using RNAscope Fluorescent Multiplex Detection Reagents (323110, Advanced Cell Diagnostics) according to the manufacturer’s instructions. Sections were then incubated with secondary antibodies (goat anti-rabbit 488, 1:500 alone; or goat anti-rabbit 488, 1:500 and goat anti-mouse 633, 1:500) at room temperature for 2 h, stained with DAPI, and mounted. Images were acquired using a Zeiss 800 confocal microscope, and the proportions of RNAscope probe-positive signals in EGFP-positive cells were quantified using ImageJ.

### Slice physiology

#### Preparation of acute brain slices

Acute brain slices were prepared following our previous study^59^. Mice were deeply anesthetized with isoflurane and decapitated, and brains were rapidly removed into ice-cold slicing solution oxygenated with 95% O_2_ and 5% CO_2_. The slicing solution contained 195 mM sucrose, 5 mM glucose, 10 mM NaCl, 25 mM NaHCO_3_, 2.5 mM KCl, 1.25 mM NaH_2_PO_4_, 2 mM Na-pyruvate, 0.5 mM CaCl_2_, and 7 mM MgCl_2_ (pH 7.3). Coronal brain slices (400 μm) were prepared using a vibratome (VT 1200S, Leica Microsystems, Wetzlar, Germany). Slices were first incubated in oxygenated artificial cerebrospinal fluid (ACSF) at 28°C for 30 min and then maintained in ACSF at room temperature for an additional 30 min before transfer to the recording chamber. ACSF contained 126 mM NaCl, 2.5 mM KCl, 2.4 mM CaCl_2_, 1.2 mM NaH_2_PO_4_, 1.2 mM MgCl_2_, 5.0 mM glucose, and 21.4 mM NaHCO_3_ (osmolarity ∼305 mOsm, pH 7.4). During recording, slices were perfused with ACSF at 1 mL/min. Acute brain slices were observed using a 40× Olympus water-immersion objective, a differential interference contrast (DIC) optical system (Olympus Inc., Japan), and a CCD camera.

#### Patch-clamp whole-cell recordings

Recording electrodes were pulled from borosilicate glass capillaries (Cat #64-0793, Warner Instruments, USA) using a PC-10 puller (Narishige Inc., Tokyo, Japan). For current-clamp recordings of action potentials, the intracellular solution contained 128 mM K-gluconate, 10 mM KCl, 10 mM HEPES, 0.1 mM EGTA, 2 mM MgCl_2_, 0.05 mM Na-GTP, and 0.05 mM Mg-ATP (pH 7.3). For voltage-clamp recordings of postsynaptic currents, the intracellular solution contained 135 mM CsCl, 10 mM HEPES, 1 mM EGTA, 2 mM QX-314 chloride, 1 mM Na-GTP, and 4 mM Mg-ATP (pH 7.4). Electrode resistance was 3.0–3.5 MΩ. Current and voltage signals were recorded using a MultiClamp 700B amplifier and Clampex 10 data acquisition software (Molecular Devices). After formation of the whole-cell recording configuration and equilibration of the intracellular solution with the cytoplasm, series resistance was compensated to 10–15 MΩ. Recordings with series resistance greater than 15 MΩ were discarded.

#### Light-evoked postsynaptic current recordings

For optical activation of channelrhodopsin-2 (ChR2)-expressing axon terminals, light pulses were delivered through a 200-μm optical fiber with the optical fiber tip positioned 500 μm above the brain slice. Laser power was adjusted to 2 mW before each experiment. Light stimulation onset, duration, and frequency were controlled by a programmable pulse generator connected to the laser system. In the presence of 4-aminopyridine (4-AP, 20 μM) and tetrodotoxin (TTX, 1 μM), light-evoked postsynaptic currents from ChR2-expressing TH-GFP+ neurons were induced by a single light pulse (2 ms) or a train of light pulses (473 nm, 2 ms, 10 Hz, 2 mW). To determine neurotransmitter receptor contributions to optically evoked postsynaptic currents, ACSF containing D-AP5 (50 μM), CNQX (20 μM), PTX (50 μM) was perfused separately.

### Chemogenetic and optogenetic manipulations

#### Chemogenetic inactivation

For chemogenetic manipulation of the LC, AAV-DIO-hM4Di-mCherry was bilaterally injected into the LC of Dbh-Cre mice, and EEG electrodes were implanted during the same procedure. For chemogenetic manipulation of LC-projecting NTS neurons, AAV2-retro-Cre was bilaterally injected into the LC of wild-type mice, and AAV-DIO-hM4Di-mCherry was bilaterally injected into the NTS, EEG electrodes were implanted during the same procedure. Behavioral and EEG testing was performed 21 days post-injection. During testing, mice were connected to the EEG recording system, and baseline activity during wakefulness was recorded. Mice then received an intraperitoneal injection of CNO (1 mg/kg). After 30 min, mice received an intraperitoneal injection of insulin (12 U/kg) or were exposed to 40% CO_2_ (for 30 min), and EEG recording continued until recovery of consciousness.

#### Optogenetic stimulation

AAV-ChR2-mCherry was bilaterally injected into the NTS of wild-type mice, optical fibers were bilaterally implanted into the LC, and EEG electrodes were implanted during the same procedure. Testing was performed 28 days post-injection. During testing, EEG activity was recorded continuously. Insulin (12 U/kg) was administered by intraperitoneal injection to induce coma, followed by optical stimulation. Single-train photostimulation (5 mW, 20 Hz, 10 s) was used to test whether activation of the NTS-LC pathway altered EEG activity and NE release. Repeated photostimulation (5 mW, 20 Hz, 10 s/min, 2 h) was used to test whether repeated activation of the NTS-LC pathway promoted recovery of consciousness after coma.

### Vagus nerve stimulation (VNS)

#### Implantation of vagus nerve cuff electrodes

Vagal electrical stimulation was performed with reference to previous research^49^. Mice were anesthetized with tribromoethanol (250 mg/kg) via intraperitoneal injection, placed in a supine position, and prepared by shaving and disinfecting the cervical surgical area with povidone-iodine. A small incision was made near the ventral midline of the neck to expose the submandibular gland and connective tissue covering the left cervical vagus nerve. The surrounding tissue was gently retracted to identify and isolate the left cervical vagus nerve while avoiding damage to the common carotid artery and internal jugular vein. The vagus nerve was gently wrapped into the cuff electrode (KedouBC, Inner diameter 0.2 mm, Interval 0.5 mm), with close contact but without visible nerve compression. A loose suture was placed between the sternohyoid and sternocleidomastoid muscles to help stabilize the cuff electrode and reduce displacement risk. The electrode lead was passed through a subcutaneous tunnel to a scalp incision. The electrode connector was secured above the cerebellum with dental cement. The cervical skin incision was sutured and coated with erythromycin ointment to prevent infection. Mice were tested after 7 days of recovery. During testing, the head-mounted connector was connected through a connecting wire to a programmable stimulator (KD-SS4, KedouBC, China), which was controlled by custom-made software through a Bluetooth-connected computer.

#### Isolated VNS combined with mesoscopic imaging

Vagus nerve cuff electrodes were implanted in Snap25-Cre::RCL-jRGECO1a-2A-NE2m mice. After 7 days, mice were fasted for 24 h, and insulin (12 U/kg) was administered by intraperitoneal injection to induce coma. After mice entered a coma-like state, short-term VNS was delivered, while mesoscopic imaging of the cerebral cortex was performed to record GRAB-NE2m signals across cortical regions. Brief stimulation was delivered with the following parameters: current intensity 1 mA, pulse width 200 μs, frequency 20 Hz, and stimulation duration 10 s.

#### Isolated VNS combined with fiber photometry

AAV-hSyn-GRAB-NE2m was unilaterally injected into the PrL of wild-type mice, and an optical fiber was implanted during the same procedure. After 21 days, a vagus nerve cuff electrode was implanted. After 7 days of recovery, mice were fasted for 24 h, and insulin (12 U/kg) was administered by intraperitoneal injection to induce coma. After mice entered a coma-like state, short-term VNS was delivered, and GRAB-NE2m fiber photometry signals and EEG signals were recorded before and after stimulation. Brief stimulation was delivered with the following parameters: current intensity 800 μA, pulse width 200 μs, frequency 20 Hz, and stimulation duration 10 s.

#### Repeated VNS to promote post-coma recovery

Vagus nerve cuff electrodes were implanted in wild-type mice. Seven days later, hypoglycemic coma was induced by insulin injection, or hypercapnic coma was induced by exposure to 40% CO_2_. After mice entered a coma-like state, repeated VNS was delivered with the following parameters: current intensity 800 μA, pulse width 200 μs, frequency 20 Hz, stimulation mode 10 s per min (10 s on / 50 s off), duration 2 h (total of 120 cycles). Control mice underwent sham vagus nerve cuff electrode implantation without VNS.

### Cell-counting strategies

Cell-counting strategies are summarized in Supplementary Table 2. For LC cell counting, 40-μm coronal sections from bregma −5.3 to −5.7 were collected from each mouse. For NTS cell counting, 40-μm coronal sections from bregma −6.24 to −8.24 were collected from each mouse. Five evenly spaced (200 μm) sections were sampled for immunohistochemistry or RNAscope labeling of marker-positive cells. Fluorescent images of the LC or NTS were acquired using an Olympus VS120 microscope with a 10× objective, and cells were counted in ImageJ using the particle analysis algorithm. Relative fluorescence intensity of tdTomato+ cell bodies or axonal terminals was measured using ImageJ. Cell counting was performed by an investigator blinded to experimental conditions.

### Quantification and statistical analysis

All experiments were performed using anonymized samples, and experimenters were blinded to experimental conditions of the mice. Statistical analyses were performed using Student *t*-test or one-way analysis of variance (ANOVA), with *n* representing the number of animals or cells. Detailed information on statistical analyses is provided in the figure legends and Supplementary Table 3. All statistical comparisons were conducted using data from at least three biologically independent experimental replicates. All data are shown as mean ± standard error of the mean (SEM).

## Reporting summary

Further information on research design is available in the Nature Portfolio Reporting Summary linked to this article.

## Data availability

All data supporting the findings of this study are provided within the paper and its Supplementary Information. All additional information will be made available upon reasonable request to the authors. Source data are provided with this paper.

## Acknowledgements

We thank Dr. Yulong Li for providing mouse lines. This work was supported by funding from NIBS and the New Cornerstone Science Foundation through the XPLORER PRIZE (P.C.). All data are archived in NIBS.

## Author contributions

Conceptualization, P.C., Z.L., F.Z., H.H.M., and T.Z.L.; investigation, M.W., L.Z., Y.L., G.C., D.L., M.Z., H.G., H.S., Y.C., and Z.Y.; writing–original draft, P.C.; funding acquisition, P.C.; supervision, P.C.

## Additional information

**Other Supplementary Materials for this manuscript include:**

Supplementary Table 1-3

Supplementary Video 1-2

**Supplementary Table 1** Summary of Reagents, organisms, and instruments.

**Supplementary Table 2** Summary of Cell counting strategies.

**Supplementary Table 3** Summary of statistical analyses.

**Supplementary Video 1** Experimental paradigm for insulin-induced hypoglycemic coma in mice. EEG was recorded and analyzed before and after insulin injection. LORR was measured after insulin injection.

**Supplementary Video 2** Experimental paradigm for CO_2_-induced hypercapnic coma in mice. EEG was recorded and analyzed before and after CO_2_ treatment. LORR was measured after CO_2_ treatment.

**Correspondence and requests for materials** should be addressed to Dr. Peng Cao.

---

**Extended Data Fig. 1.**
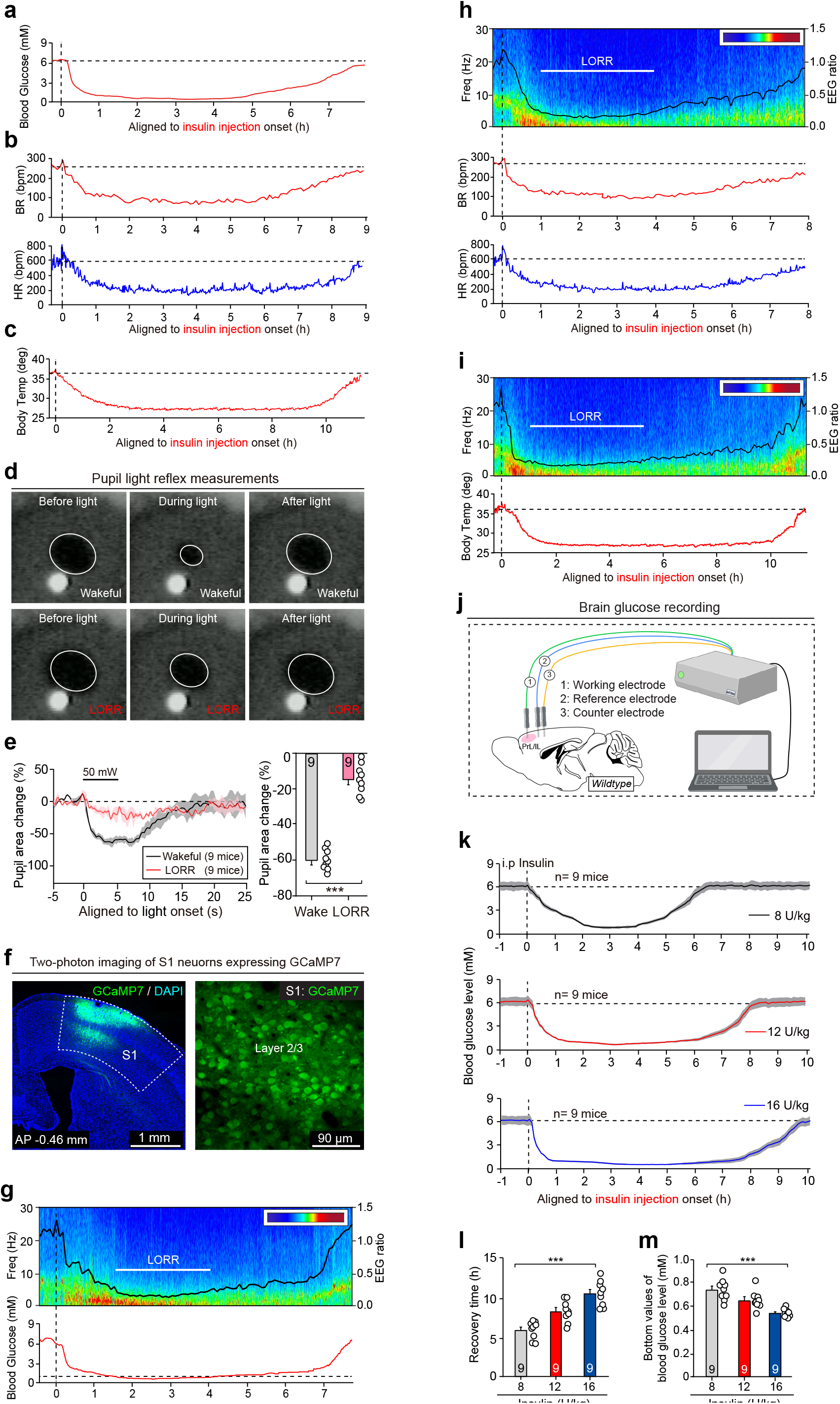
Paradigm for insulin-induced hypoglycemic coma in mice. **(a-c)** Representative traces of blood glucose level (a), breathing rate (BR) (b, *top*), heart rate (HR) (b, *bottom*), and body temperature (c) before and after intraperitoneal injection of insulin (12 U/kg) in mice. Vertical dashed lines indicate time points for insulin injection. **(d)** Representative images showing pupil size before, during, and after light illumination in a representative mouse during wakefulness and insulin-induced LORR. **(e)** Time courses (*left*) and quantification of normalized pupil area changes (*right*), showing pupil light reflex during wakefulness and LORR. **(f)** *Left*, coronal brain section showing GCaMP7 expression in S1. *Right*, representative micrograph showing GCaMP7 expression in layer 2/3 S1 neurons. **(g)** Time course of blood glucose level in parallel with insulin-induced coma state of a representative mouse. **(h)** Time courses of breathing rate (BR) and heart rate (HR) in parallel with insulin-induced coma state of a representative mouse. **(i)** Time course of body temperature in parallel with insulin-induced coma state of a representative mouse. **(j)** Schematic showing methods to measure intracerebral glucose level. **(k)** Averaged time courses of blood glucose level before and after insulin injection with different dosages (8, 12, and 16 U/kg). **(l, m)** Quantification of blood glucose recovery time (l) and minimum values of blood glucose (m) in response to insulin at different dosages (8, 12, and 16 U/kg). Scale bars are labeled in graphs. Numbers of mice (e, k-m) are indicated in graphs. Data in (e, k-m) are mean ± SEM. Statistical analyses (e, l, m) were performed using Student *t*-test (*** *P* < 0.001) or one-way ANOVA (*** *P* < 0.001). For *P*-values, see Supplementary Table 3.

**Extended Data Fig. 2.**
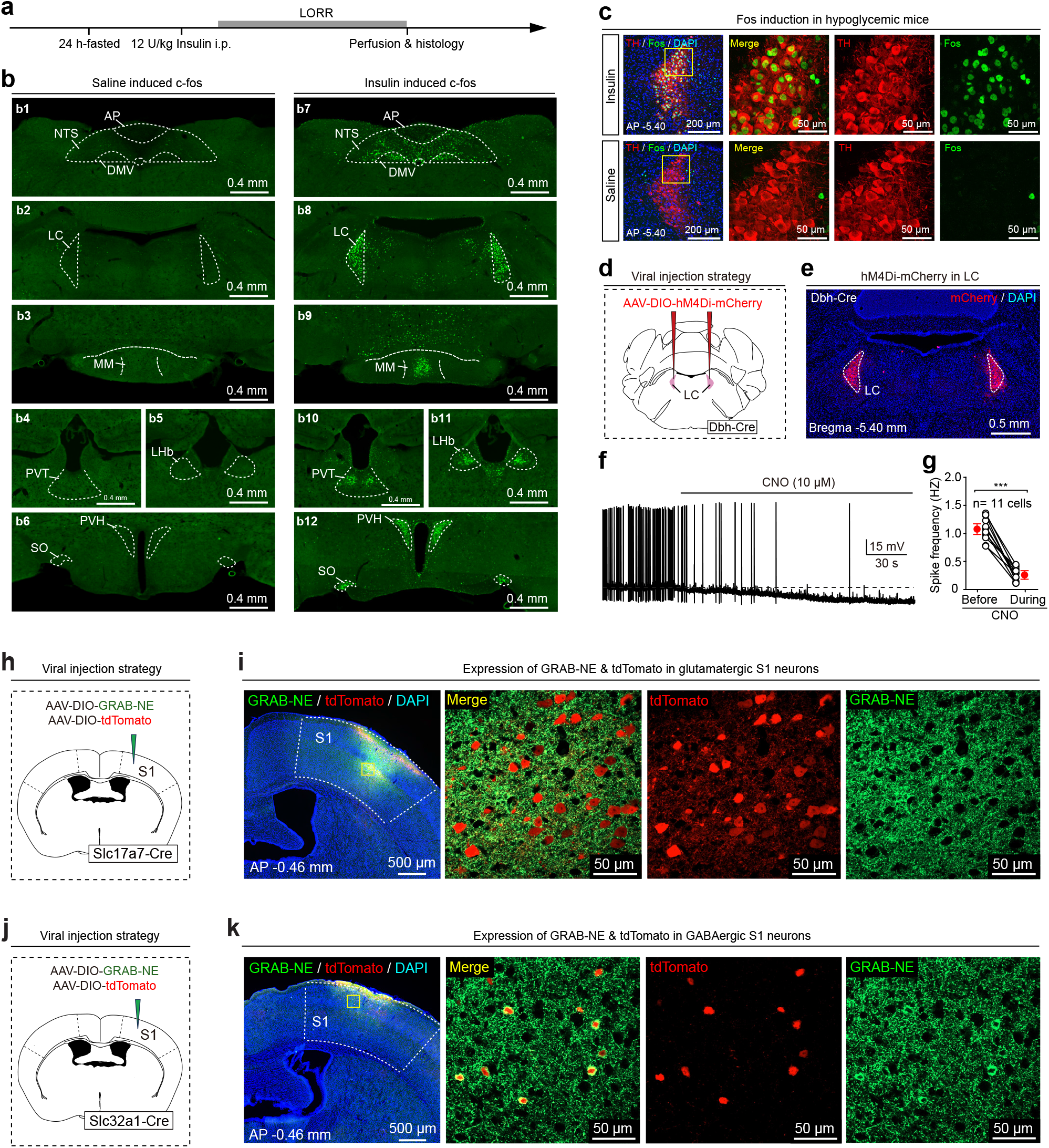
Role of LC NE neurons in insulin-induced hypoglycemic coma. **(a)** Schematic showing strategy used to measure Fos expression in mice subjected to insulin-induced hypoglycemic coma. **(b)** Representative micrographs showing Fos expression in different brain areas of mice without (b1–b6) or with (b7–b12) insulin-induced hypoglycemic coma. Abbreviations: AP, area postrema; NTS, nucleus tractus solitarius; DMV, dorsal motor nucleus of the vagus nerve; LC, locus coeruleus; MM, medial mammillary nucleus, medial part; PVT, periventricular thalamic nucleus; LHb, lateral habenula; PVH, periventricular hypothalamic nucleus; SO, supraoptic nucleus. **(c)** Representative micrographs showing Fos expression induced in LC NE neurons of mice subjected to insulin-induced hypoglycemic coma. Tyrosine hydroxylase (TH) was used as a marker of LC NE neurons. **(d)** Schematic showing bilateral injection of AAV-DIO-hM4Di-mCherry in the LC of Dbh-Cre mice. **(e)** Representative micrograph showing expression of hM4Di-mCherry in the LC. **(f, g)** Representative traces of action potential firing (f) and quantification of firing frequency (g), showing CNO perfusion in ACSF effectively reduced action potential firing in hM4Di-expressing LC NE neurons. **(h)** Schematic showing injection of AAV-DIO-GRAB-NE2m mixed with AAV-DIO-tdTomato (1:1) in S1 of Slc17a7-Cre mice. **(i)** Representative micrographs showing expression of GRAB-NE2m and tdTomato in glutamatergic S1 neurons in Slc17a7-Cre mice. **(j)** Schematic showing injection of AAV-DIO-GRAB-NE2m mixed with AAV-DIO-tdTomato (1:1) in S1 of Slc32a1-Cre mice. **(k)** Representative micrographs showing expression of GRAB-NE2m and tdTomato in GABAergic S1 neurons in Slc32a1-Cre mice. Scale bars are labeled in graphs. Numbers of cells (g) are indicated in graphs. Data in (g) are mean ± SEM. Statistical analyses (g) were performed using Student *t-*test (*** *P* < 0.001). For *P*-values, see Supplementary Table 3.

**Extended Data Fig. 3.**
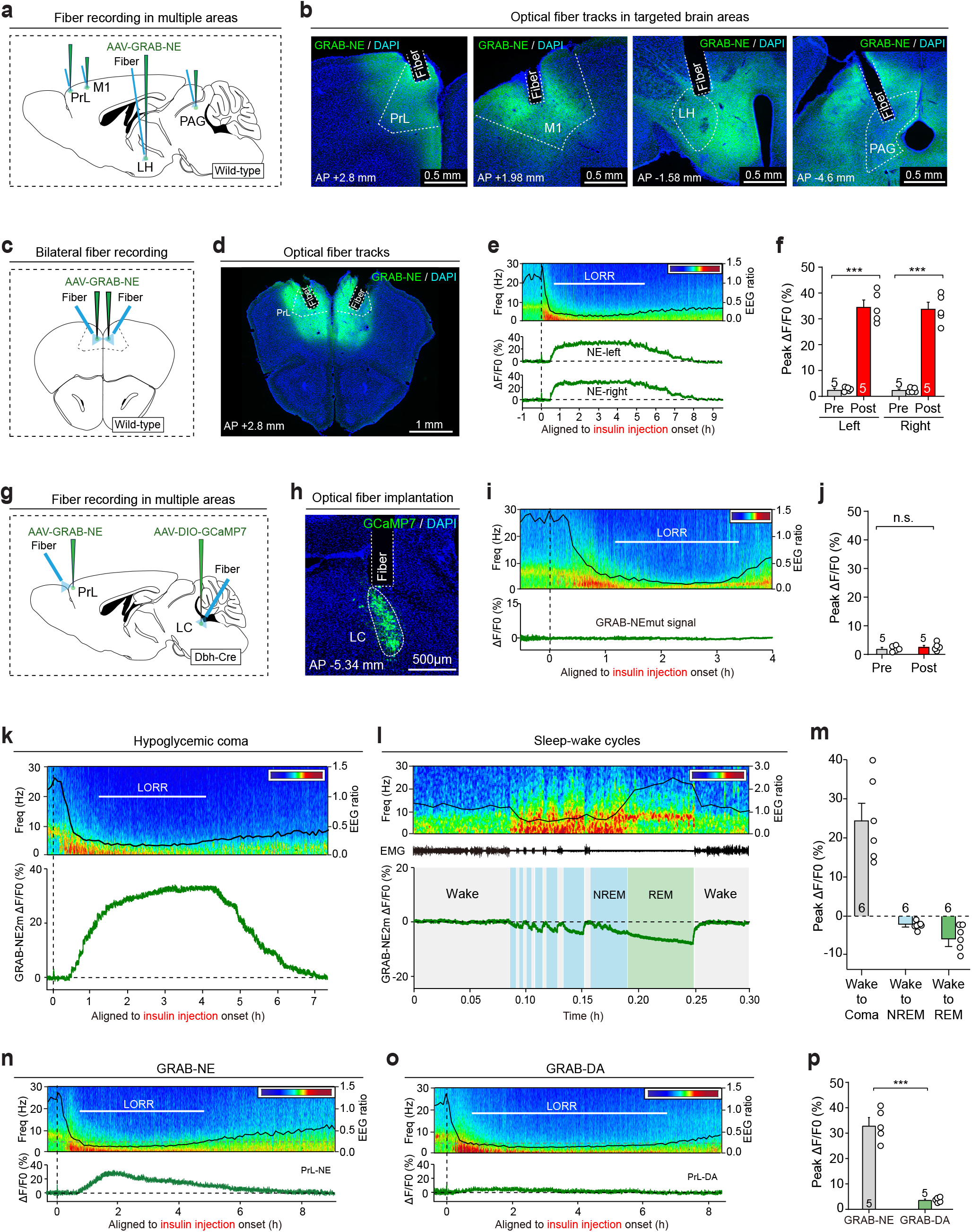
Brain-wide NE release evoked by insulin-induced hypoglycemia. **(a)** Schematic showing AAV-GRAB-NE2m injection into different brain areas and optical fiber implantation for simultaneous multisite fiber photometry recordings. **(b)** Representative micrographs showing optical fiber tracks above AAV-GRAB-NE2m-infected brain regions. **(c)** Schematic showing bilateral AAV-GRAB-NE2m injection into the PrL, followed by optical fiber implantation. **(d)** Representative micrographs showing optical fiber tracks above AAV-GRAB-NE2m-infected PrL. **(e)** Representative traces of GRAB-NE2m fluorescence changes in left and right PrL, shown with insulin-induced hypoglycemic coma state. **(f)** Quantification of peak GRAB-NE2m ΔF/F0 in bilateral PrL before and during insulin-induced hypoglycemic coma. **(g)** Schematic showing strategy used for simultaneous recording of GRAB-NE2m signals in the PrL and GCaMP signals from LC NE neurons. **(h)** Representative micrograph showing optical fiber track above GCaMP7-expressing LC NE neurons. **(i)** Representative trace of GRAB-NEmut fluorescence changes (as a control), shown with insulin-induced hypoglycemic coma state. **(j)** Quantification showing no significant changes in GRAB-NEmut fluorescence in the PrL of mice subjected to insulin-induced hypoglycemic coma. **(k)** Representative trace of GRAB-NE2m fluorescence changes, shown with insulin-induced coma state in a representative mouse. **(l)** Representative trace of GRAB-NE2m fluorescence changes, shown with brain states during multiple episodes of sleep-wake cycles. **(m)** Quantification comparing GRAB-NE2m fluorescence changes during wakefulness-to-coma transition, wakefulness-to-NREM transition, and wakefulness-to-REM transition. **(n)** Representative trace of GRAB-NE2m fluorescence changes, shown with insulin-induced coma state in a representative mouse. **(o)** Representative trace of GRAB-DA2m fluorescence changes, shown with insulin-induced coma state in a representative mouse. **(p)** Quantification of peak ΔF/F0 of GRAB-NE2m versus GRAB-DA2m in mice subjected to insulin-induced hypoglycemic coma. Scale bars are labeled in graphs. Numbers of mice (f, j, m, p) are indicated in graphs. Data in (f, j, m, p) are mean ± SEM. Statistical analyses (f, j, p) were performed using Student *t-*test (n.s. *P* > 0.1; *** *P* < 0.001). For *P*-values, see Supplementary Table 3.

**Extended Data Fig. 4.**
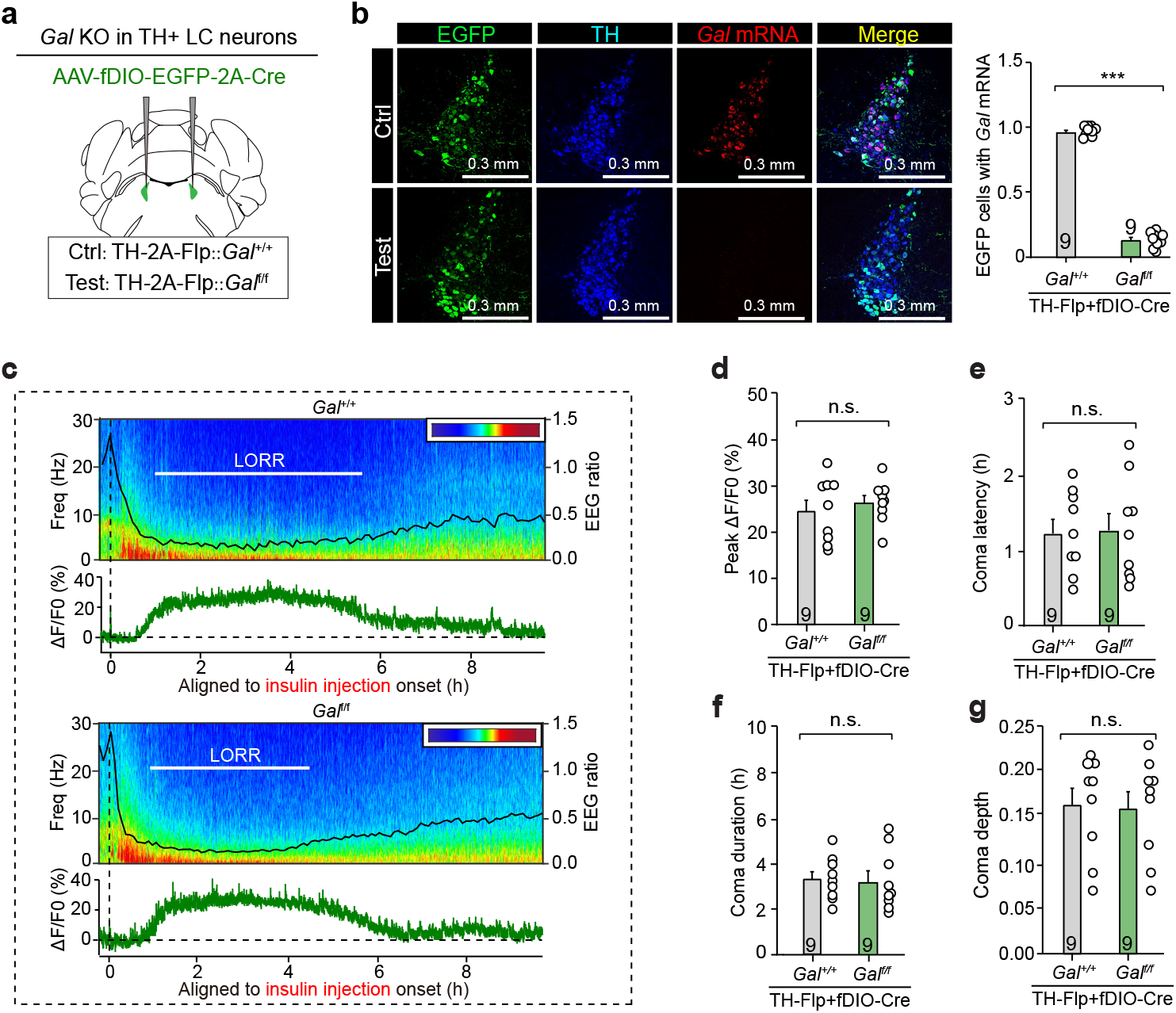
Effects of *Gal* deletion in LC NE neurons on insulin-induced hypoglycemic coma. **(a)** Schematic showing strategy used for *Gal* deletion in LC NE neurons. **(b)** Representative micrographs (*left*) and quantification (*right*) showing efficient deletion of *Gal* in LC NE neurons. **(c)** Representative traces of GRAB-NE2m fluorescence changes, shown with insulin-induced hypoglycemic coma state in mice with or without deletion of *Gal* in LC NE neurons. **(d)** Quantification of peak GRAB-NE2m ΔF/F0 in mice with or without deletion of *Gal* in LC NE neurons. **(e-g)** Quantification of insulin-induced hypoglycemic coma latency (e), duration (f), and depth (g) in mice with or without deletion of *Gal* in LC NE neurons. Numbers of mice (b, d-g) are indicated in graphs. Data in (b, d-g) are mean ± SEM. Statistical analyses (b, d-g) were performed using Student *t-*test (*** *P* < 0.001; n.s. *P* > 0.1). For *P*-values, see Supplementary Table 3.

**Extended Data Fig. 5.**
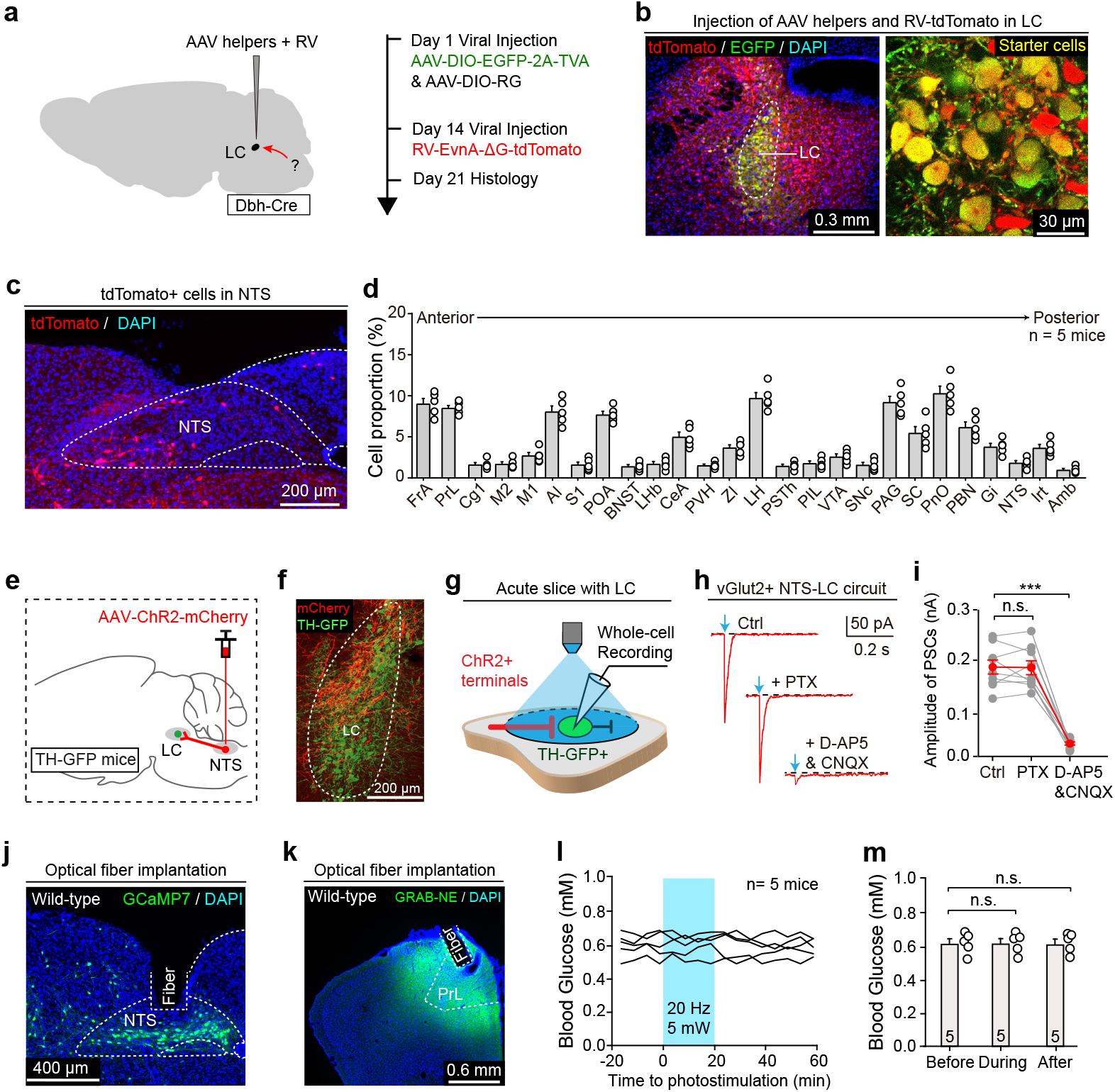
Characterization of NTS-LC circuit. **(a)** Schematic showing strategy used for retrograde labeling of afferent neurons that monosynaptically innervate LC NE neurons with recombinant G-deleted rabies virus (RV). **(b)** Representative micrographs showing labeling of starter cells in the LC. **(c)** Representative micrograph showing tdTomato+ NTS neurons labeled by RV. **(d)** Quantification of tdTomato+ neuron distribution in different brain regions. **(e)** Schematic showing injection of AAV-ChR2-mCherry in the NTS of TH-GFP mice. **(f)** Representative micrograph showing ChR2-mCherry+ axons intermingled with TH-GFP+ neurons in the LC. **(g)** Schematic showing whole-cell recording of TH-GFP+ neurons in the LC paired with photostimulation of ChR2-expressing axon terminals from NTS in the acute brain slice. **(h, i)** Representative traces of light-evoked postsynaptic currents (h) and quantification of current amplitude (i) in response to GABAa receptor antagonist (PTX) or glutamate receptor antagonists (D-AP5, CNQX). **(j)** Representative micrograph showing optical fiber track above LC-projecting NTS neurons expressing GCaMP7. **(k)** Representative micrograph showing optical fiber track above AAV-GRAB-NE2m-infected PrL. **(l, m)** Representative traces (l) and quantification (m) of blood glucose levels before, during, and after photostimulation of NTS-LC circuit in mice with hypoglycemic coma. Numbers of mice (d, m) or cells (i) are indicated in graphs. Data in (d, i, m) are mean ± SEM. Statistical analyses (i, m) were performed using Student *t-*test (*** *P* < 0.001; n.s. *P* > 0.1). For *P*-values, see Supplementary Table 3.

**Extended Data Fig. 6.**
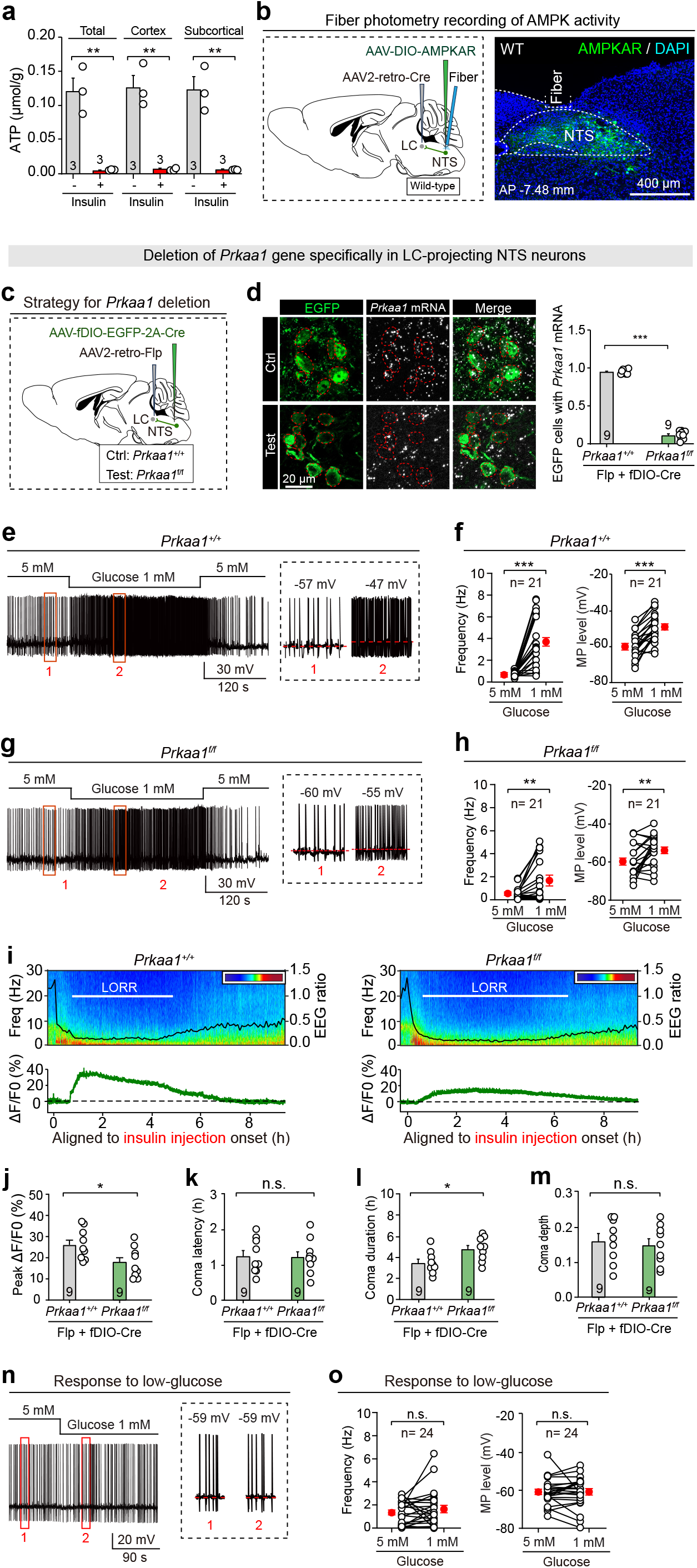
Activation of LC-projecting NTS neurons by hypoglycemia depends on AMPK signaling. **(a)** Quantification of brain ATP levels in mice with or without insulin-induced hypoglycemic coma. **(b)** *Left*, schematic showing strategy used for fiber recording of AMPKAR fluorescence in LC-projecting NTS neurons. *Right*, representative micrograph showing optical fiber track above AMPKAR-expressing LC-projecting NTS neurons. **(c)** Schematic showing strategy for *Prkaa1* deletion in LC-projecting NTS neurons. **(d)** Representative micrographs (*left*) and quantification (*right*) showing efficient deletion of *Prkaa1* in LC-projecting NTS neurons. **(e, g)** Representative traces of action potential firing (*left*) and membrane potential (*right*) in LC-projecting NTS neurons without (e) and with (g) *Prkaa1* deletion in response to low-glucose stimulation. **(f, h)** Quantification of action potential firing frequency (*left*) and membrane potential level (*right*) in LC-projecting NTS neurons without (f) and with (h) *Prkaa1* deletion in response to low-glucose stimulation. **(i)** Representative traces of GRAB-NE2m signals, shown with coma state in mice without (*left*) and with (*right*) *Prkaa1* deletion in LC-projecting NTS neurons. **(j)** Quantification of GRAB-NE2m signals, showing effects of *Prkaa1* deletion in LC-projecting NTS neurons on cortical NE release after hypoglycemia. **(k-m)** Quantification of effects of *Prkaa1* deletion in LC-projecting NTS neurons on hypoglycemic coma latency (k), duration (l), and depth (m). **(n)** Representative traces of action potential firing (*left*) and membrane potential (*right*) showing low-glucose responses of an LC NE neuron in TH-GFP mice. **(o)** Quantification of action potential firing frequency (*left*) and membrane potential level (*right*) in all recorded LC NE neurons to low-glucose stimulation. Numbers of mice (a, d, j-m) or cells (f, h, o) are indicated in graphs. Data in (a, d, f, h, j-m, o) are mean ± SEM. Statistical analyses (a, d, f, h, j-m, o) were performed using Student *t-*test (*** *P* < 0.001; ** *P* < 0.01; * *P* < 0.05; n.s. *P* > 0.1). For *P*-values, see Supplementary Table 3.

**Extended Data Fig. 7.**
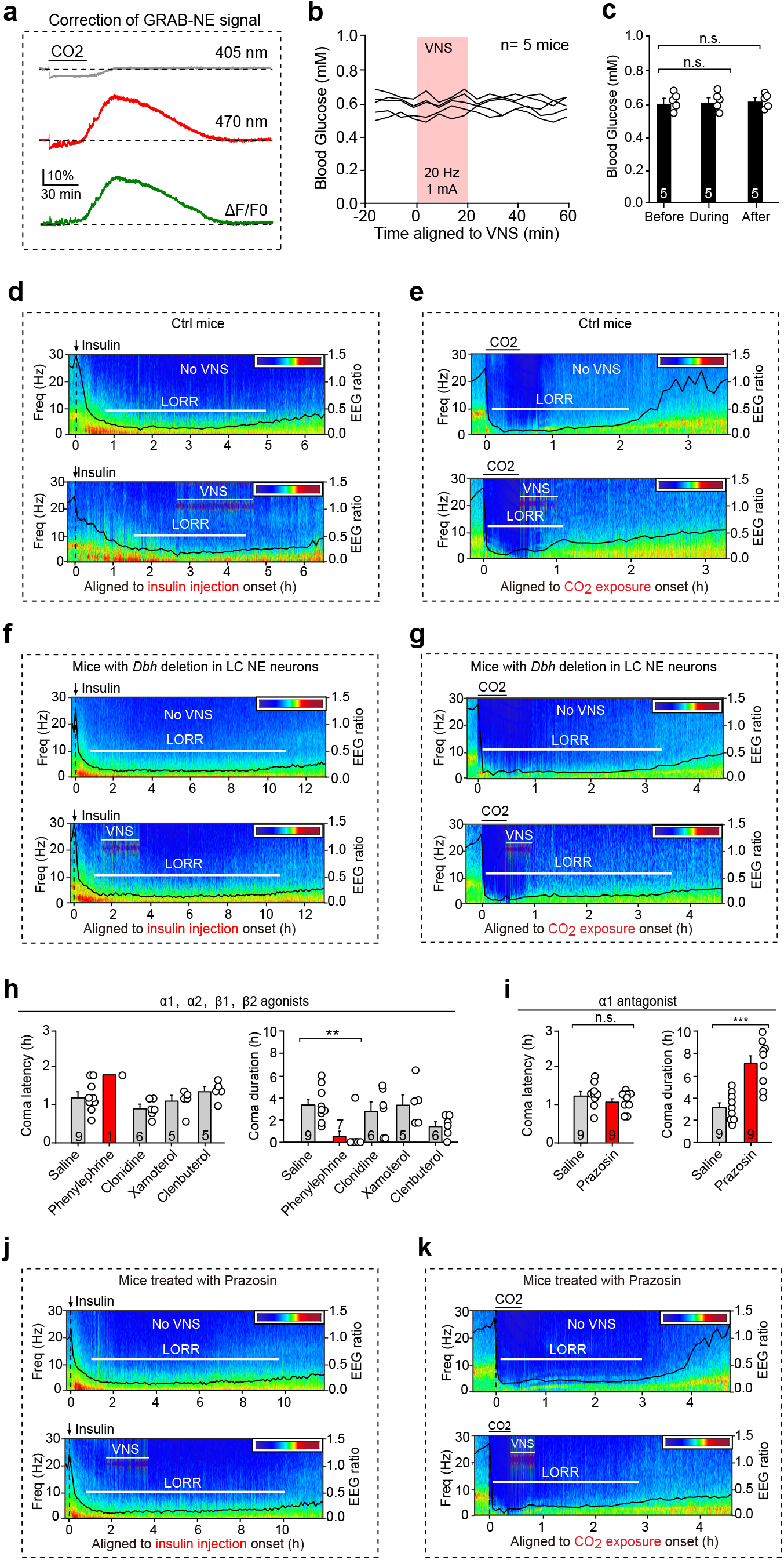
Vagus nerve stimulation (VNS) evokes NE release critical for post-coma arousal. **(a)** Schematic showing correction procedure used to compensate for CO_2_-related artifacts in GRAB-NE2m fluorescence signals. See “Correction of NE signals during hypercapnia” in Methods Details. **(b, c)** Representative traces (b) and quantification (c) of blood glucose level before, during, and after VNS. **(d, e)** Representative EEG heat-maps, EEG ratio curves and LORR timing showing effect of VNS on insulin-induced hypoglycemic coma (d) or CO_2_-induced hypercapnic coma (e) in wild-type mice. **(f, g)** Representative EEG heat-maps, EEG ratio curves and LORR timing showing effect of VNS on insulin-induced hypoglycemic coma (f) or CO_2_-induced hypercapnic coma (g) in mice with *Dbh* deletion in LC NE neurons. **(h)** Quantification of coma latency (*left*) and duration (*right*), showing effects of agonists targeting different adrenergic receptors (α1, phenylephrine, 10 mg/kg; α2, clonidine, 2 mg/kg; β1, xamoterol, 3 mg/kg; β2, clenbuterol, 5 mg/kg) on insulin-induced hypoglycemic coma in wild-type mice. **(i)** Quantification of coma latency (*left*) and duration (*right*), showing effects of α1 adrenergic receptor antagonist (prazosin, 6 mg/kg) on insulin-induced hypoglycemic coma in wild-type mice. **(j, k)** Representative EEG heat-maps, EEG ratio curves and LORR timing showing effect of VNS on insulin-induced hypoglycemic coma (j) or CO_2_-induced hypercapnic coma (k) in mice pretreated with prazosin. Numbers of mice (c, h, i) are indicated in graphs. Data in (c, h, i) are mean ± SEM. Statistical analyses (c, h, i) were performed using Student *t-*test (*** *P* < 0.001; ** *P* < 0.01; n.s. *P* > 0.1). For *P*-values, see Supplementary Table 3.

