## Supplementary Table 1 for "AMPK-dependent brain-wide norepinephrine release promotes recovery from metabolic coma in mice"

### Supplementary Table 1 Summary of reagents, organisms, and instruments

| Reagents, Organisms or instruments | Company | Catalog Number |
|---|---|---|
| Antibodies | | |
| Rabbit polyclonal anti-EGFP | Abcam | Cat # ab290 |
| Chicken polyclonal anti-EGFP | Abcam | Cat # ab13970 |
| Chicken polyclonal anti-mCherry | Abcam | Cat# ab205402 |
| Rabbit monoclonal anti-c-Fos | Cell Signaling Technology | Cat # 2250s |
| Rabbit polyclonal anti-TH | Merck Millipore | Cat # AB152 |
| Mouse monoclonal anti-TH | Immunostar | Cat # 22941 |
| Rabbit polyclonal anti-Dbh | Invitrogen | Cat # PA5-34664 |
| Bacterial and virus strains | | |
| AAV2/9-hSyn-jGCaMP7f | TaiTool Co., Ltd | N/A |
| AAV2/9-hSyn-DIO-jGCaMP7f | TaiTool Co., Ltd | N/A |
| AAV2/9-hSyn-DIO-AMPKAR | TaiTool Co., Ltd | N/A |
| AAV2/9-hSyn-DIO-AMPKAR T/A | TaiTool Co., Ltd | N/A |
| rAAV-hSyn-NE2m | BrainCase Co., Ltd | N/A |
| rAAV-hSyn-NE2m-mut | BrainCase Co., Ltd | N/A |
| rAAV-hSyn-DA2m | BrainCase Co., Ltd | N/A |
| rAAV-hSyn-DIO-NE2m | BrainCase Co., Ltd | N/A |
| rAAV-EF1α-DIO-tdTomato | BrainCase Co., Ltd | N/A |
| AAV2/9-hSyn-DIO-mCherry | TaiTool Co., Ltd | N/A |
| AAV2/9-hSyn-DIO-hM4Di-mCherry | TaiTool Co., Ltd | N/A |
| AAV2-retro-hSyn-Cre | TaiTool Co., Ltd | N/A |
| AAV2-retro-hSyn-Flp | TaiTool Co., Ltd | N/A |
| AAV2/9-EF1α-fDIO-EGFP-2A-Cre | TaiTool Co., Ltd | N/A |
| AAV2/9-hSyn-mCherry | TaiTool Co., Ltd | N/A |
| AAV2/9-hSyn-ChR2-mCherry | TaiTool Co., Ltd | N/A |
| scAAV2/1-hSyn-Cre | TaiTool Co., Ltd | N/A |
| AAV2/9-hSyn-DIO-EYFP | TaiTool Co., Ltd | N/A |
| AAV2/9-EF1α-DIO-EGFP-2A-TVA | BrainVTA Co., Ltd. | N/A |
| AAV2/9-EF1α-DIO-RVG | BrainVTA Co., Ltd. | N/A |
| RV-EnvA-ΔG-tdTomato | BrainVTA Co., Ltd. | N/A |
| Chemicals, peptides, and recombinant proteins | | |
| Insulin | NovoRapid | N/A |
| 40% CO2 | Linde | N/A |
| CNO | Enzo Life Sciences | Cat # BML-NS105 |
| DAPI | Sigma | Cat # D8417 |
| D-AP5 | Tocris | Cat # 0106 |
| CNQX | Tocris | Cat # 0190 |
| PTX | Tocris | Cat # 3097 |
| Compound C | MedChemExpress | Cat # HY-13418A |
| AICAR | MedChemExpress | Cat # HY-13417 |

| | | |
|---|---|---|
| Phenylephrine hydrochloride | MedChemExpress | Cat # HY-B0471 |
| Prazosin hydrochloride | Sigma | Cat # P7791 |
| Clonidine hydrochloride | Sigma | Cat # C7897 |
| Xamoterol hemifumarate | MedChemExpress | Cat # HY-101327A |
| Clenbuterol hydrochloride | MedChemExpress | Cat # HY-B1614 |
| RNAscope Probe-Mm-Gal | Advanced Cell Diagnostics | Cat # 400961 |
| RNAscope Probe-Mm-Prkaa1-O1 | Advanced Cell Diagnostics | Cat # 430831 |
| RNAscope Probe-Mm-Prkaa2-O1 | Advanced Cell Diagnostics | Cat # 430841 |
| Critical commercial assays | | |
| RNAscope Multiplex Fluorescent Reagent Kit | Advanced Cell Diagnostics | Cat # 323110 |
| Enhanced ATP Assay Kit | Beyotime | Cat # S0027 |
| Experimental models: Organisms/strains | | |
| Snap25-Cre | Jackson Laboratory | Cat # JAX: 023525 |
| RCL-jRGECO1a-2A-NE2m | Yulong Li Lab | N/A |
| Dbh-Cre | MMRRC | Cat # 032081-UCD |
| Slc17a7-Cre | Jackson Laboratory | Cat # JAX: 023527 |
| Slc32a1-Cre | Jackson Laboratory | Cat # JAX: 016962 |
| TH-2A-Flp | MMRRC | Cat # 050618-MU |
| Ai14 | Jackson Laboratory | Cat # JAX: 007914 |
| TH-GFP | Jackson Laboratory | Cat # RBRC03162 |
| Dbh f/f | GemPharmatech | Cat # T017958 |
| Gal f/f | GemPharmatech | Cat # T013199 |
| Prkaa1 f/f | GemPharmatech | Cat # T007182 |
| Prkaa2 f/f | GemPharmatech | Cat # T013159 |
| Software and algorithms | | |
| LabChart | ADInstruments | https://www.adinstruments.com.cn/ |
| MATLAB | Mathworks | https://www.mathworks.com |
| ImageJ | NIH | https://imagej.nih.gov/ij/index.html |
| Ponemah | Data Sciences International | https://www.datasci.com |
| FinePointe | Data Sciences International | https://www.datasci.com |
| AnalyzIR | FOTRIC | https://www.fotric.cn |
| Mosaic 3.0 | TUCSEN | https://www.tucsen.com |
| pClamp 10 | Molecular Devices | https://www.moleculardevices.com/ |
| Other | | |
| PowerLab | ADInstruments | https://www.adinstruments.com.cn/ |
| DSI MX2 | Data Sciences International | https://www.datasci.com |
| DSI Buxco | Data Sciences International | https://www.datasci.com |
| FOTRIC 226s | FOTRIC | https://www.fotric.cn |
| MT200 | MT Technology | https://www.mingtaijiaxin.com |

| | | |
|---|---|---|
| KD-SS4 | KedouBC | https://www.kedoubc.com |
| Olympus VS120 | Olympus | https://www.olympus-lifescience.com |
| Carl Zeiss LSM 800 | Carl Zeiss | https://www.zeiss.com |
| Nikon A1+ MP | Nikon | https://www.microscope.healthcare.nikon.com |
| Olympus MVX10 | Olympus | https://www.olympus-lifescience.com |
| MultiClamp 700B amplifier | Molecular Devices | https://www.moleculardevices.com/ |
