## Supplementary Table 2 for "AMPK-dependent brain-wide norepinephrine release promotes recovery from metabolic coma in mice"

### Supplementary Table 2 Summary of cell-counting strategies

| Brain region | Section Type | Section Range | Total collection | Sampling |
|---|---|---|---|---|
| LC | Coronal section (40 μm) | Bregma (-5.3 to -5.7) | Approximately 10 sections | To sample all the sections |
| NTS | Coronal section (40 μm) | Bregma (-6.24 to -8.24) | Approximately 50 sections | To sample all the sections |
| FrA | Coronal section (40 μm) | Bregma (+3.7 to +2.2) | Approximately 38 sections | To sample all the sections |
| PrL | Coronal section (40 μm) | Bregma (+3.1 to +1.5) | Approximately 40 sections | To sample all the sections |
| IL | Coronal section (40 μm) | Bregma (+2.0 to +1.2) | Approximately 20 sections | To sample all the sections |
| Cg1 | Coronal section (40 μm) | Bregma (+2.4 to -0.3) | Approximately 68 sections | To sample all the sections |
| AuD | Coronal section (40 μm) | Bregma (-2.0 to -3.8) | Approximately 45 sections | To sample all the sections |
| V1 | Coronal section (40 μm) | Bregma (-2.1 to -5.0) | Approximately 73 sections | To sample all the sections |
| S1 | Coronal section (40 μm) | Bregma (+2.0 to -2.3) | Approximately 108 sections | To sample all the sections |
| M2 | Coronal section (40 μm) | Bregma (+2.5 to -1.4) | Approximately 98 sections | To sample all the sections |
| M1 | Coronal section (40 μm) | Bregma (+2.5 to -1.4) | Approximately 98 sections | To sample all the sections |
| CeA | Coronal section (40 μm) | Bregma (-0.8 to -1.8) | Approximately 25 sections | To sample all the sections |
| AI | Coronal section (40 μm) | Bregma (+2.5 to -1.3) | Approximately 95 sections | To sample all the sections |
| POA | Coronal section (40 μm) | Bregma (+0.7 to -0.5) | Approximately 30 sections | To sample all the sections |
| MPO | Coronal section (40 μm) | Bregma (+0.7 to -0.5) | Approximately 30 sections | To sample all the sections |
| BNST | Coronal section (40 μm) | Bregma (+0.6 to -0.6) | Approximately 30 sections | To sample all the sections |
| LHb | Coronal section (40 μm) | Bregma (-0.9 to -2.3) | Approximately 35 sections | To sample all the sections |
| PVT | Coronal section (40 μm) | Bregma (-0.2 to -2.3) | Approximately 52 sections | To sample all the sections |
| SuM | Coronal section (40 μm) | Bregma (-2.4 to -3.2) | Approximately 20 sections | To sample all the sections |
| MM | Coronal section (40 μm) | Bregma (-2.7 to -3.3) | Approximately 15 sections | To sample all the sections |

| | | | | |
|---|---|---|---|---|
| SO | Coronal section (40 μm) | Bregma (-0.5 to -1.5) | Approximately 25 sections | To sample all the sections |
| PVH | Coronal section (40 μm) | Bregma (-0.6 to -1.2) | Approximately 15 sections | To sample all the sections |
| ZI | Coronal section (40 μm) | Bregma (-0.9 to -2.9) | Approximately 50 sections | To sample all the sections |
| LH | Coronal section (40 μm) | Bregma (-0.3 to -2.8) | Approximately 60 sections | To sample all the sections |
| PH | Coronal section (40 μm) | Bregma (-1.8 to -2.7) | Approximately 23 sections | To sample all the sections |
| PSTh | Coronal section (40 μm) | Bregma (-2.1 to -2.5) | Approximately 10 sections | To sample all the sections |
| PIL | Coronal section (40 μm) | Bregma (-2.7 to -3.4) | Approximately 18 sections | To sample all the sections |
| VTA | Coronal section (40 μm) | Bregma (-2.8 to -3.8) | Approximately 25 sections | To sample all the sections |
| SNc | Coronal section (40 μm) | Bregma (-2.8 to -3.8) | Approximately 25 sections | To sample all the sections |
| PAG | Coronal section (40 μm) | Bregma (-3.0 to -5.0) | Approximately 50 sections | To sample all the sections |
| SG | Coronal section (40 μm) | Bregma (-3.0 to -3.8) | Approximately 20 sections | To sample all the sections |
| SC | Coronal section (40 μm) | Bregma (-3.3 to -4.6) | Approximately 33 sections | To sample all the sections |
| EW | Coronal section (40 μm) | Bregma (-2.8 to -4.0) | Approximately 30 sections | To sample all the sections |
| DRV | Coronal section (40 μm) | Bregma (-4.0 to -5.2) | Approximately 30 sections | To sample all the sections |
| PnO | Coronal section (40 μm) | Bregma (-4.0 to -5.0) | Approximately 25 sections | To sample all the sections |
| PBN | Coronal section (40 μm) | Bregma (-5.0 to -5.7) | Approximately 18 sections | To sample all the sections |
| Nucleus O | Coronal section (40 μm) | Bregma (-5.3 to -5.7) | Approximately 10 sections | To sample all the sections |
| Gi | Coronal section (40 μm) | Bregma (-5.5 to -7.1) | Approximately 40 sections | To sample all the sections |
| LDTg | Coronal section (40 μm) | Bregma (-4.7 to -5.4) | Approximately 18 sections | To sample all the sections |
| IO | Coronal section (40 μm) | Bregma (-6.7 to -8.0) | Approximately 33 sections | To sample all the sections |
| DMV | Coronal section (40 μm) | Bregma (-6.4 to -8.1) | Approximately 42 sections | To sample all the sections |
| LRT | Coronal section (40 μm) | Bregma (-7.0 to -8.2) | Approximately 30 sections | To sample all the sections |

| | | | | |
|---|---|---|---|---|
| IRT | Coronal section (40 μm) | Bregma (-5.4 to -8.2) | Approximately 70 sections | To sample all the sections |
| Amb | Coronal section (40 μm) | Bregma (-6.6 to -8.2) | Approximately 40 sections | To sample all the sections |
| NG/JG | Horizontal section (40 μm) | From dorsal to ventral | Approximately 26 sections | To sample all the sections |
