## Supplementary Table 3 for "AMPK-dependent brain-wide norepinephrine release promotes recovery from metabolic coma in mice"

### Supplementary Table 3 Summary of statistical analyses

| Figure | Sample size (n) | Statistical test | P values |
|---|---|---|---|
| Fig. 1d | Before = 20 cells | Student t-test | Wakeful During vs. Before: P= 2.30128E-06 *** |
| | During = 20 cells | | LORR During vs. Before: P= 0.3299 n.s. |
| Fig. 1e | Wakeful = 20 cells | Student t-test | LORR vs. Wakeful: P= 3.46898-06 *** |
| | LORR = 20 cells | | |
| Fig. 1f | Wakeful = 6 mice | Student t-test | LORR vs. Wakeful: P= 5.27904E-06 *** |
| | LORR = 6 mice | | |
| Fig. 1i | Wake = 15 mice | Student t-test | LORR vs. Wake: P= 1.18154E-11 *** |
| | NREM = 15 mice | | LORR vs. NREM: P= 3.08323E-9 *** |
| | REM = 15 mice | | LORR vs. REM: P= 9.62986E-12 *** |
| | LORR = 15 mice | | |
| Fig. 1l | 8 U/kg = 9 mice | One-Way ANOVA | 16 U/kg vs. 12 U/kg vs. 8 U/kg: P= 1.61E-05 *** |
| | 12 U/kg = 9 mice | | |
| | 16 U/kg = 9 mice | | |
| Fig. 1m | 8 U/kg = 9 mice | One-Way ANOVA | 16 U/kg vs. 12 U/kg vs. 8 U/kg: P= 3.57E-05 *** |
| | 12 U/kg = 9 mice | | |
| | 16 U/kg = 9 mice | | |
| Fig. 1n | 8 U/kg = 9 mice | One-Way ANOVA | 16 U/kg vs. 12 U/kg vs. 8 U/kg: P= 9.78E-06 *** |
| | 12 U/kg = 9 mice | | |
| | 16 U/kg = 9 mice | | |
| Fig. 1s | Ctrl = 9 mice | Student t-test | hM4Di vs. Ctrl: P= 0.7304 n.s. |
| | hM4Di = 9 mice | | |
| Fig. 1t | Ctrl = 9 mice | Student t-test | hM4Di vs. Ctrl: P= 0.0013 ** |
| | hM4Di = 9 mice | | |
| Fig. 1u | Ctrl = 9 mice | Student t-test | hM4Di vs. Ctrl: P= 0.0303 * |
| | hM4Di = 9 mice | | |
| Fig. 2d | Pre = 5 mice | Student t-test | M1 Post vs. Pre: P= 3.625E-06 *** |
| | Post = 5 mice | | S1 Post vs. Pre: P= 4.556E-06 *** |
| | | | V1 Post vs. Pre: P= 4.898E-06 *** |
| Fig. 2f | Before = 7 mice | Student t-test | During vs. Before: P= 0.0002 *** |
| | During = 7 mice | | |
| Fig. 2g | Before = 7 mice | Student t-test | During vs. Before: P= 0.0019 ** |
| | During = 7 mice | | |
| Fig. 2j | Pre = 5 mice | Student t-test | PrL Post vs. Pre: P= 0.0009 *** |
| | Post = 5 mice | | M1 Post vs. Pre: P= 0.0008 *** |
| | | | LH Post vs. Pre: P= 0.0007 *** |
| | | | PAG Post vs. Pre: P= 0.0006 *** |
| Fig. 2k | Pre = 5 mice | Student t-test | LC-GCaMP Post vs. Pre: P= 0.0051 ** |
| | Post = 5 mice | | PrL-NE Post vs. Pre: P= 0.0006 *** |
| Fig. 2m | *Dbh*$^{+/+}$ = 9 mice | Student t-test | *Dbh*$^{f/f}$ vs. *Dbh*$^{+/+}$: P= 1.6E-13 *** |
| | *Dbh*$^{f/f}$ = 9 mice | | |
| Fig. 2o | *Dbh*$^{+/+}$ = 9 mice | Student t-test | *Dbh*$^{f/f}$ vs. *Dbh*$^{+/+}$: P= 0.0002 *** |

| | | | |
|---|---|---|---|
| | *Dbh*$^{f/f}$ = 9 mice | | |
| Fig. 2p | *Dbh*$^{+/+}$ = 9 mice<br>*Dbh*$^{f/f}$ = 9 mice | Student t-test | *Dbh*$^{f/f}$ vs. *Dbh*$^{+/+}$: P= 0.6309 n.s. |
| Fig. 2q | *Dbh*$^{+/+}$ = 9 mice<br>*Dbh*$^{f/f}$ = 9 mice | Student t-test | *Dbh*$^{f/f}$ vs. *Dbh*$^{+/+}$: P= 5.64996E-07 *** |
| Fig. 2r | *Dbh*$^{+/+}$ = 9 mice<br>*Dbh*$^{f/f}$ = 9 mice | Student t-test | *Dbh*$^{f/f}$ vs. *Dbh*$^{+/+}$: P= 0.0461 * |
| Fig. 3f | Pre = 5 mice<br>Post = 5 mice | Student t-test | NTS-GCaMP Post vs. Pre: P= 0.0023 **<br>PrL-NE Post vs. Pre: P= 0.0002 *** |
| Fig. 3i | Before = 9 mice<br>During = 9 mice | Student t-test | During vs. Before: P= 2.05088E-07 *** |
| Fig. 3j | Before = 9 mice<br>During = 9 mice | Student t-test | During vs. Before: P= 8.42957E-06 *** |
| Fig. 3l | Ctrl = 9 mice<br>ChR2 = 9 mice | Student t-test | ChR2 vs. Ctrl: P= 0.0054 ** |
| Fig. 3m | Ctrl = 9 mice<br>ChR2 = 9 mice | Student t-test | ChR2 vs. Ctrl: P= 0.011 * |
| Fig. 3q | Ctrl = 9 mice<br>hM4Di = 9 mice | Student t-test | hM4Di vs. Ctrl: P= 2.36394E-05 *** |
| Fig. 3r | Ctrl = 9 mice<br>hM4Di = 9 mice | Student t-test | hM4Di vs. Ctrl: P= 0.6102 n.s. |
| Fig. 3s | Ctrl = 9 mice<br>hM4Di = 9 mice | Student t-test | hM4Di vs. Ctrl: P= 2.27749E-05 *** |
| Fig. 3t | Ctrl = 9 mice<br>hM4Di = 9 mice | Student t-test | hM4Di vs. Ctrl: P= 0.0476 * |
| Fig. 4b | 5mM = 21 cells<br>1mM = 21 cells | Student t-test | Frequency 1mM vs. 5mM: P= 1.37249E-06 ***<br>MP 1mM vs. 5mM: P= 3.474E-06 *** |
| Fig. 4e | 5mM = 21 cells<br>1mM = 21 cells | Student t-test | Frequency 1mM vs. 5mM: P= 3.79127E-05 ***<br>MP 1mM vs. 5mM: P= 6.02305E-06 *** |
| Fig. 4g | 5mM = 21 cells<br>1mM = 21cells | Student t-test | Frequency 1mM vs. 5mM: P= 0.0533 n.s.<br>MP 1mM vs. 5mM: P= 0.9123 n.s. |
| Fig. 4i | Before = 21 cells<br>During = 21 cells | Student t-test | Frequency During vs. Before: P= 2.24662E-06 ***<br>MP During vs. Before: P= 3.439E-09 *** |
| Fig. 4k | Before = 21 cells<br>During = 21 cells | Student t-test | Frequency During vs. Before: P= 0.174 n.s.<br>MP During vs. Before: P= 0.13 n.s. |
| Fig. 4l | T/A-mut = 9 mice<br>ExRai = 9 mice | Student t-test | ExRai vs. T/A-mut: P= 3.3269E-05 *** |
| Fig. 4n | *Prkaa2*$^{+/+}$ = 9 mice<br>*Prkaa2*$^{f/f}$ = 9 mice | Student t-test | *Prkaa2*$^{f/f}$ vs. *Prkaa2*$^{+/+}$: P= 2.639E-11 *** |
| Fig. 4p | 5mM = 21 cells<br>1mM = 21 cells | Student t-test | Frequency 1mM vs. 5mM: P= 4.73479E-06 ***<br>MP 1mM vs. 5mM: P= 2.022E-07 *** |
| Fig. 4r | 5mM = 21 cells<br>1mM = 21 cells | Student t-test | Frequency 1mM vs. 5mM: P= 0.2834 n.s.<br>MP 1mM vs. 5mM: P= 0.5881 n.s. |

| | | | |
|---|---|---|---|
| Fig. 4t | *Prkaa2*$^{+/+}$ = 9 mice<br>*Prkaa2*$^{f/f}$ = 9 mice | Student t-test | *Prkaa2*$^{f/f}$ vs. *Prkaa2*$^{+/+}$: P= 0.0001 *** |
| Fig. 4u | *Prkaa2*$^{+/+}$ = 9 mice<br>*Prkaa2*$^{f/f}$ = 9 mice | Student t-test | Coma latency *Prkaa2*$^{f/f}$ vs. *Prkaa2*$^{+/+}$: P= 0.7791 n.s.<br>Coma duration *Prkaa2*$^{f/f}$ vs. *Prkaa2*$^{+/+}$: P= 6.71593E-05 ***<br>Coma depth *Prkaa2*$^{f/f}$ vs. *Prkaa2*$^{+/+}$: P= 0.0478 * |
| Fig. 5c | Wake = 15 mice<br>NREM = 15 mice<br>REM = 15 mice<br>LORR = 15 mice | Student t-test | LORR vs. Wake: P= 4.20838E-10 ***<br>LORR vs. NREM: P= 6.23524E-09 ***<br>LORR vs. REM: P= 3.65144E-12 *** |
| Fig. 5d | Total = 3 mice<br>CTX = 3 mice<br>Sub-CTX = 3 mice | Student t-test | Total CO2 + vs. -: P= 0.0153 *<br>CTX CO2 + vs. -: P= 0.0157 *<br>Sub-CTX CO2 + vs. -: P= 0.012 * |
| Fig. 5e | T/A-mut = 9 mice<br>ExRai = 9 mice | Student t-test | ExRai vs. T/A-mut: P= 2.09214E-06 *** |
| Fig. 5f | Pre = 9 mice<br>Post = 9 mice | Student t-test | Post vs.Pre: P= 1.83291E-07 *** |
| Fig. 5h | *Prkaa2*$^{+/+}$ = 9 mice<br>*Prkaa2*$^{f/f}$ = 9 mice | Student t-test | *Prkaa2*$^{f/f}$ vs. *Prkaa2*$^{+/+}$: P= 3.2774E-07 *** |
| Fig. 5i | *Prkaa2*$^{+/+}$ = 9 mice<br>*Prkaa2*$^{f/f}$ = 9 mice | Student t-test | Coma latency *Prkaa2*$^{f/f}$ vs. *Prkaa2*$^{+/+}$: P= 0.6561 n.s.<br>Coma duration *Prkaa2*$^{f/f}$ vs. *Prkaa2*$^{+/+}$: P= 0.0004 ***<br>Coma depth *Prkaa2*$^{f/f}$ vs. *Prkaa2*$^{+/+}$: P= 0.0345 * |
| Fig. 5k | *Dbh*$^{+/+}$ = 9 mice<br>*Dbh*$^{f/f}$ = 9 mice | Student t-test | *Dbh*$^{f/f}$ vs. *Dbh*$^{+/+}$: P= 1.23901E-06 *** |
| Fig. 5l | *Dbh*$^{+/+}$ = 9 mice<br>*Dbh*$^{f/f}$ = 9 mice | Student t-test | Coma latency *Dbh*$^{f/f}$ vs. *Dbh*$^{+/+}$: P= 0.7275 n.s.<br>Coma duration *Dbh*$^{f/f}$ vs. *Dbh*$^{+/+}$: P= 0.0006 ***<br>Coma depth *Dbh*$^{f/f}$ vs. *Dbh*$^{+/+}$: P= 0.0407 * |
| Fig. 6f | Before = 5 mice<br>During = 5 mice | Student t-test | M1 During vs. Before: P= 0.0001 ***<br>S1 During vs. Before: P= 0.0003 ***<br>V1 During vs. Before: P= 0.0006 *** |
| Fig. 6h | Before = 9 mice<br>During = 9 mice | Student t-test | During vs. Before: P= 2.82917E-07 *** |
| Fig. 6i | Before = 9 mice<br>During = 9 mice | Student t-test | During vs. Before: P= 2.57113E-06 *** |
| Fig 6k | Before = 9 mice<br>During = 9 mice | Student t-test | During vs. Before: P= 5.08654E-08 *** |
| Fig. 6l | Before = 9 mice<br>During = 9 mice | Student t-test | During vs. Before: P= 8.38709E-06 *** |
| Fig. 6m | Sham = 9 mice<br>VNS= 9 mice | Student t-test | Coma latency VNS vs. Sham: P= 0.8669 n.s.<br>Coma duration VNS vs. Sham: P= 0.0294 *<br>Coma depth VNS vs. Sham: P= 0.0266 * |
| Fig. 6n | Sham = 9 mice<br>VNS= 9 mice | Student t-test | Coma latency VNS vs. Sham: P= 0.5685 n.s.<br>Coma duration VNS vs. Sham: P= 0.0016 **<br>Coma depth VNS vs. Sham: P= 0.0481 * |

| | | | |
|---|---|---|---|
| Fig. 6o | Sham = 9 mice<br>VNS= 9 mice | Student t-test | Coma latency VNS vs. Sham: P= 0.9094 n.s.<br>Coma duration VNS vs. Sham: P= 0.5936 n.s.<br>Coma depth VNS vs. Sham: P= 0.9371 n.s. |
| Fig. 6p | Sham = 9 mice<br>VNS= 9 mice | Student t-test | Coma latency VNS vs. Sham: P= 0.8978 n.s.<br>Coma duration VNS vs. Sham: P= 0.9635 n.s.<br>Coma depth VNS vs. Sham: P= 0.7906 n.s. |
| Fig. 6q | Sham = 9 mice<br>VNS= 9 mice | Student t-test | Coma latency VNS vs. Sham: P= 0.9631 n.s.<br>Coma duration VNS vs. Sham: P= 0.6815 n.s.<br>Coma depth VNS vs. Sham: P= 0.7932 n.s. |
| Fig. 6r | Sham = 9 mice<br>VNS= 9 mice | Student t-test | Coma latency VNS vs. Sham: P= 0.6691 n.s.<br>Coma duration VNS vs. Sham: P= 0.7244 n.s.<br>Coma depth VNS vs. Sham: P= 0.9497 n.s. |
| Extended Data Fig. 1e | Wake = 9 mice<br>LORR = 9 mice | Student t-test | LORR vs. Wake: P= 1.94626E-06 *** |
| Extended Data Fig. 1l | 8 U/kg = 9 mice<br>12 U/kg = 9 mice<br>16 U/kg = 9 mice | One-Way ANOVA | 16 U/kg vs. 12 U/kg vs. 8 U/kg: P= 1.23E-06 *** |
| Extended Data Fig. 1m | 8 U/kg = 9 mice<br>12 U/kg = 9 mice<br>16 U/kg = 9 mice | One-Way ANOVA | 16 U/kg vs. 12 U/kg vs. 8 U/kg: P= 2.75E-05 *** |
| Extended Data Fig. 2g | Before = 11 cells<br>During = 11 cells | Student t-test | During vs. Before: P= 2.83642E-07 *** |
| Extended Data Fig. 3f | Pre = 5 mice<br>Post = 5 mice | Student t-test | Left Post vs. Pre: P= 0.0002 ***<br>Right Post vs. Pre: P= 0.0002 *** |
| Extended Data Fig. 3j | Pre = 5 mice<br>Post = 5 mice | Student t-test | Post vs. Pre: P= 0.5454 n.s. |
| Extended Data Fig. 3p | GRAB-NE = 5 mice<br>GRAB-DA = 5 mice | Student t-test | GRAB-DA vs. GRAB-NE: P= 0.0004 *** |
| Extended Data Fig. 4b | *Gal* $^{+/+}$ = 9 mice<br>*Gal* $^{f/f}$ = 9 mice | Student t-test | *Gal* $^{f/f}$ vs. *Gal* $^{+/+}$: P= 8.10284E-15 *** |
| Extended Data Fig. 4d | *Gal* $^{+/+}$ = 9 mice<br>*Gal* $^{f/f}$ = 9 mice | Student t-test | *Gal* $^{f/f}$ vs. *Gal* $^{+/+}$: P= 0.5254 n.s. |
| Extended Data Fig. 4e | *Gal* $^{+/+}$ = 9 mice<br>*Gal* $^{f/f}$ = 9 mice | Student t-test | *Gal* $^{f/f}$ vs. *Gal* $^{+/+}$: P= 0.9397 n.s. |
| Extended Data Fig. 4f | *Gal* $^{+/+}$ = 9 mice<br>Gal $^{f/f}$ = 9 mice | Student t-test | *Gal* $^{f/f}$ vs. *Gal* $^{+/+}$: P= 0.8737 n.s. |
| Extended Data Fig. 4g | *Gal* $^{+/+}$ = 9 mice<br>*Gal* $^{f/f}$ = 9 mice | Student t-test | *Gal* $^{f/f}$ vs. *Gal* $^{+/+}$: P= 0.8994 n.s. |
| Extended Data Fig. 5i | Ctrl = 9 mice<br>+PTX = 9 mice<br>+D-AP5&CNQX = 9 mice | Student t-test | +PTX vs. Ctrl: P= 0.81 n.s.<br>+D-AP5&CNQX vs. Ctrl: P= 5.74871E-06 *** |
| Extended Data Fig. 5m | Before = 5 mice<br>During = 5 mice | Student t-test | During vs. Before: P= 0.8276 n.s.<br>After vs. Before: P= 0.8544 n.s. |

| | | | |
|---|---|---|---|
| | After = 5 mice | | |
| Extended Data Fig. 6a | Total = 3 mice<br>Cortex = 3 mice<br>Subcortical = 3 mice | Student t-test | Total Insulin + vs. -: P= 0.0044 **<br>Cortex Insulin + vs. -: P= 0.0025 **<br>Subcortical Insulin + vs. -: P= 0.0036 ** |
| Extended Data Fig. 6d | *Prkaa1*$^{+/+}$ = 9 mice<br>*Prkaa1*$^{f/f}$ = 9 mice | Student t-test | *Prkaa1*$^{f/f}$ vs. *Prkaa1*$^{+/+}$: P= 9.4129E-14 *** |
| Extended Data Fig. 6f | 5mM = 21 cells<br>1mM = 21 cells | Student t-test | Frequency 1mM vs. 5mM: P= 8.4542E-06 ***<br>MP 1mM vs. 5mM: P= 3.2935E-07 *** |
| Extended Data Fig. 6h | 5mM = 21 cells<br>1mM = 21 cells | Student t-test | Frequency 1mM vs. 5mM: P= 0.0034 **<br>MP 1mM vs. 5mM: P= 0.0022 ** |
| Extended Data Fig. 6j | *Prkaa1*$^{+/+}$ = 9 mice<br>*Prkaa1*$^{f/f}$ = 9 mice | Student t-test | *Prkaa1*$^{f/f}$ vs. *Prkaa1*$^{+/+}$: P= 0.0237 * |
| Extended Data Fig. 6k | *Prkaa1*$^{+/+}$ = 9 mice<br>*Prkaa1*$^{f/f}$ = 9 mice | Student t-test | *Prkaa1*$^{f/f}$ vs. *Prkaa1*$^{+/+}$: P= 0.9634 n.s. |
| Extended Data Fig. 6l | *Prkaa1*$^{+/+}$ = 9 mice<br>*Prkaa1*$^{f/f}$ = 9 mice | Student t-test | *Prkaa1*$^{f/f}$ vs. *Prkaa1*$^{+/+}$: P= 0.0213 * |
| Extended Data Fig. 6m | *Prkaa1*$^{+/+}$ = 9 mice<br>*Prkaa1*$^{f/f}$ = 9 mice | Student t-test | *Prkaa1*$^{f/f}$ vs. *Prkaa1*$^{+/+}$: P= 0.6427 n.s. |
| Extended Data Fig. 6o | 5mM = 24 cells<br>1mM = 24 cells | Student t-test | Frequency 1mM vs. 5mM: P= 0.3854 n.s.<br>MP 1mM vs. 5mM: P= 0.9468 n.s. |
| Extended Data Fig. 7c | Before = 5 mice<br>During = 5 mice<br>After = 5 mice | Student t-test | During vs. Before: P= 0.4766 n.s.<br>After vs. Before: P= 0.5122 n.s. |
| Extended Data Fig. 7h | Saline = 9 mice<br>Phenylephrine = 7 mice | Student t-test | Coma duration Phenylephrine vs. Saline: P= 0.003 ** |
| Extended Data Fig. 7i | Saline = 9 mice<br>Prazosin = 9 mice | Student t-test | Coma latency Prazosin vs. Saline: P= 0.1896 n.s.<br>Coma duration Prazosin vs. Saline: P= 0.0001 *** |
